# Whole-genome duplication drives biosynthetic gene cluster fragmentation and regulatory rewiring of monoterpene indole alkaloid metabolism in *Strychnos*

**DOI:** 10.64898/2026.08.19.745744

**Authors:** Jiming Liu, Joan Jing Yi Jong, Rami-Petteri Apuli, Hebi Zhuang, Roy Jun Kai Tham, Abner Herbert Lim, Wei Liu, Jia Jun Ngiam, Matti A. Niissalo, Gillian S. Khew, Bin Tean Teh, Jarkko Salojärvi

## Abstract

Whole-genome duplications (WGDs) reshape plant genomes by generating redundancy, after which lineage-specific architectures emerge through fractionation, gene loss and rearrangement. How specialized metabolic pathways remain functionally integrated after such large-scale restructuring remains poorly understood. This problem is especially relevant for biosynthetic gene clusters (BGCs), which physically organize specialized-metabolism genes yet can be disrupted by post-duplication rearrangement. Here, we present the first chromosome-level genomes for Loganiaceae, including near telomere-to-telomere assemblies of *Strychnos ignatii* and *S. pubescens*, together with a draft genome of the extinct species *S. ridleyi*. Following a lineage-specific WGD, the two extant *Strychnos* species evolved contrasting genome-evolutionary trajectories and metabolite profiles: *S. ignatii* shows expansion of monoterpenoid- and monoterpene indole alkaloid (MIA)-associated gene families and strychnine-type MIA dominance, whereas *S. pubescens* exhibits elevated transposable element activity associated with DNA-binding with one finger (DOF)-linked regulatory rewiring and broader sesquiterpenoid- and triterpenoid-rich chemistry. Crucially, both species retain active strychnine biosynthesis despite fragmentation of a deeply conserved alkaloid BGC in MIA-producing Gentianales, revealing how pathway function can persist after disruption of ancestral BGC architecture. Comparative metabolomic and transcriptomic pathway analyses indicate norfluorocurarine oxidase (*NO*) as a major divergence point associated with strychnine accumulation. Promoter analyses, yeast one-hybrid assays, and electrophoretic mobility shift assays support a model in which *S. ignatii* retains the canonical jasmonate-responsive *MYB*, *MYC2/bHLH*, and *AP2/ERF cis*-regulatory module at *NO*, whereas the orthologous *S. pubescens* promoter shows reduced capacity to recruit these activators and instead exhibits a DOF-associated architecture. Together, our results show that WGD can decouple physical cluster architecture from pathway function, allowing specialized metabolic pathways to remain active while divergent chemical phenotypes evolve through lineage-specific combinations of coding-space expansion and transposable-element-associated *cis*-regulatory rewiring.

## Introduction

Whole-genome duplications (WGDs) are among the most important drivers of plant genome evolution, generating extensive genetic redundancy that can facilitate innovation in morphology, physiology, and specialized metabolism^1^. Following WGD, duplicated genes and genomic regions undergo fractionation through differential loss, rearrangement and regulatory divergence, often resulting in substantial restructuring of metabolic pathways. Biosynthetic gene clusters (BGCs), genomic regions containing physically linked genes that contribute to specialized metabolism, represent one class of genomic architecture that may be particularly sensitive to these processes. Such clusters, including the Bx1–Bx5 cluster for DIBOA biosynthesis in maize^2^, the avenacin cluster in oat, the noscapine cluster in *Papaver somniferum*, and the thalianol cluster in *Arabidopsis*^3^, are thought to facilitate coordinated regulation, inheritance and maintenance of critical enzymatic steps. Post-duplication restructuring may therefore affect specialized metabolism both by changing enzymatic coding space through differential gene retention and duplication, and by reshaping regulatory sequence space through rearrangements and transposable-element activity. How metabolic pathways maintain functional integration while generating metabolic diversity during such large-scale genome restructuring remains poorly understood.

Monoterpene indole alkaloids (MIAs), produced by plants in the Gentianales, Garryales and Cornales, provide an attractive system for investigating these questions. With more than 3,000 compounds identified to date^4^, MIAs are among the most structurally diverse and functionally important classes of plant specialized metabolites, ranging from the anticancer drugs vincristine and vinblastine to the potent neurotoxin strychnine^5,6^. Their exceptional diversity makes MIAs a powerful system for understanding how genome evolution contributes to metabolic diversification and innovation in plants. The order Gentianales is a major source of MIA biosynthetic diversity and includes many medicinally important plants across its five families: Apocynaceae, Gelsemiaceae, Gentianaceae, Loganiaceae, and Rubiaceae. Within Apocynaceae, *Catharanthus roseus* serves as a model species whose anticancer alkaloid pathway has been extensively characterized^7^, whereas *Gelsemium sempervirens* represents Gelsemiaceae with diverse MIAs^8^. In Rubiaceae, *Ophiorrhiza pumila* is a key system for studying camptothecin biosynthesis^9^. By contrast, MIA biosynthesis appears to have been lost in Gentianaceae^10^, while Loganiaceae is still largely lacking genomic studies, despite the recent elucidation of the strychnine biosynthesis pathway^11^.

Comparative genomic analyses across these representative Gentianales species have shown that genes involved in synthesis of the MIA precursor strictosidine, namely tryptophan decarboxylase (*TDC*), strictosidine synthase (*STR*), and multidrug and toxic compound extrusion (*MATE*) transporter genes, are physically linked in a conserved BGC. This *STR–TDC–MATE* cluster is thought to have originated near the base of Gentianales and represents one of the most evolutionarily conserved genomic architectures associated with plant specialized metabolism. Functionally, the *STR*-*TDC*-*MATE* cluster generates strictosidine, the universal precursor of all MIAs. *TDC* produces tryptamine, which is then transported into the vacuole by *MATE*^12,13^. There, *STR* condenses the monoamine with secologanin, an iridoid derived from the broader terpenoid network, in a Pictet-Spengler reaction to form strictosidine. Despite this conserved entry point, downstream metabolic profiles differ substantially among species, reflecting extensive diversification of MIA pathways across Gentianales. Furthermore, because MIA biosynthesis relies on carbon flux from upstream terpenoid pathways, genomic restructuring may not only alter MIA diversity but also dictate whether pathway output is biased toward MIAs or alternative terpenoid defenses. The *STR–TDC–MATE* cluster is not an isolated structural arrangement, as a recent study identified two additional BGCs contributing to MIA biosynthesis in Rubiaceae and Apocynaceae^14^, underscoring that physical linkage is a recurring evolutionary strategy for managing these complex pathways.

Despite increasing genetic resources in Gentianales, Loganiaceae has remained absent from genomic studies of MIA evolution. This omission is particularly striking because the family occupies a key phylogenetic position for reconstructing the evolutionary history of MIA biosynthetic pathways. This pantropical family consists of approximately 460 species in 16 genera, of which *Strychnos* L. is the largest, comprising 200 species^15^. *Strychnos* is best known for strychnine, a highly toxic corynanthe-type MIA that accumulates mainly in seeds and roots^16,17^. However, the genus also exhibits substantial metabolic diversity. For example, evolutionary divergence of an acetyltransferase (*AT*) enzyme redirects the canonical strychnine pathway towards diaboline in *Strychnos potatorum*, whereas strychnine remains the predominant MIA in *S. nux-vomica*^11^. Such variation at a low taxonomic level indicates recent evolutionary shifts in pathway activity and metabolite output, making *Strychnos* an attractive genus for investigating how specialized metabolic pathways diversify through genome evolution and regulatory change.

Here, using PacBio HiFi and Omni-C sequencing, we generated the first high-quality near telomere-to-telomere (T2T) reference genome assemblies of *Strychnos ignatii* and *S. pubescens* from Singapore, and reconstructed the genome of the now extinct *S. ridleyi* from a 19th-century specimen^18^. By integrating comparative genomics, transcriptomics, metabolomics and regulatory analyses, we investigate how whole-genome duplication, fragmentation of ancestral BGCs, and *cis*-regulatory evolution have shaped diversification of MIA metabolism in *Strychnos*. Our analyses reveal that active strychnine biosynthesis has been maintained despite fragmentation of the canonical *STR–TDC–MATE* cluster following lineage-specific WGD, and show that divergent chemical phenotypes evolved through contrasting combinations of coding-space expansion and transposable-element-associated *cis*-regulatory rewiring. These findings provide a framework for understanding how post-duplication genome restructuring can decouple physical cluster architecture from pathway function while generating metabolic diversity in plants.

### Near chromosome-complete *Strychnos* genome assemblies provide a foundation for studying MIA pathway evolution

To establish a genomic foundation for investigating specialized metabolism in Loganiaceae, we generated the first high-quality, chromosome-level reference genomes for *S. ignatii* and *S. pubescens*. Utilizing PacBio HiFi long-read sequencing (yielding ∼80x coverage) and Omni-C scaffolding, we generated highly contiguous chromosome-scale assemblies approaching telomere-to-telomere (T2T) contiguity (**Supplementary Table S1**). The final assemblies anchored 99.4% and 97.8% of the assembled sequence to 20 pseudochromosomes in *S. ignatii* and *S. pubescens*, respectively, leaving only 17 unplaced scaffolds in each assembly (**Fig. 1A, B; Supplementary Figs. S1-S5, Supplementary Tables S1-S4**). The assemblies also captured 36 telomeric and 20 centromeric regions in each species. The gene-coding space was highly complete, with the *S. ignatii* and *S. pubescens* assemblies recovering 98.2% and 97.9% of the universally conserved orthologs (BUSCO v5.4.2 with eudicot ODB10 dataset; **Fig. 1D**), respectively. Despite the shared, highly contiguous karyotype, the two genomes exhibited marked structural divergence, with the *S. pubescens* assembly (660 Mb) being substantially larger than that of *S. ignatii* (480 Mb**; Supplementary Tables S2–S3**).

**Figure 1.**
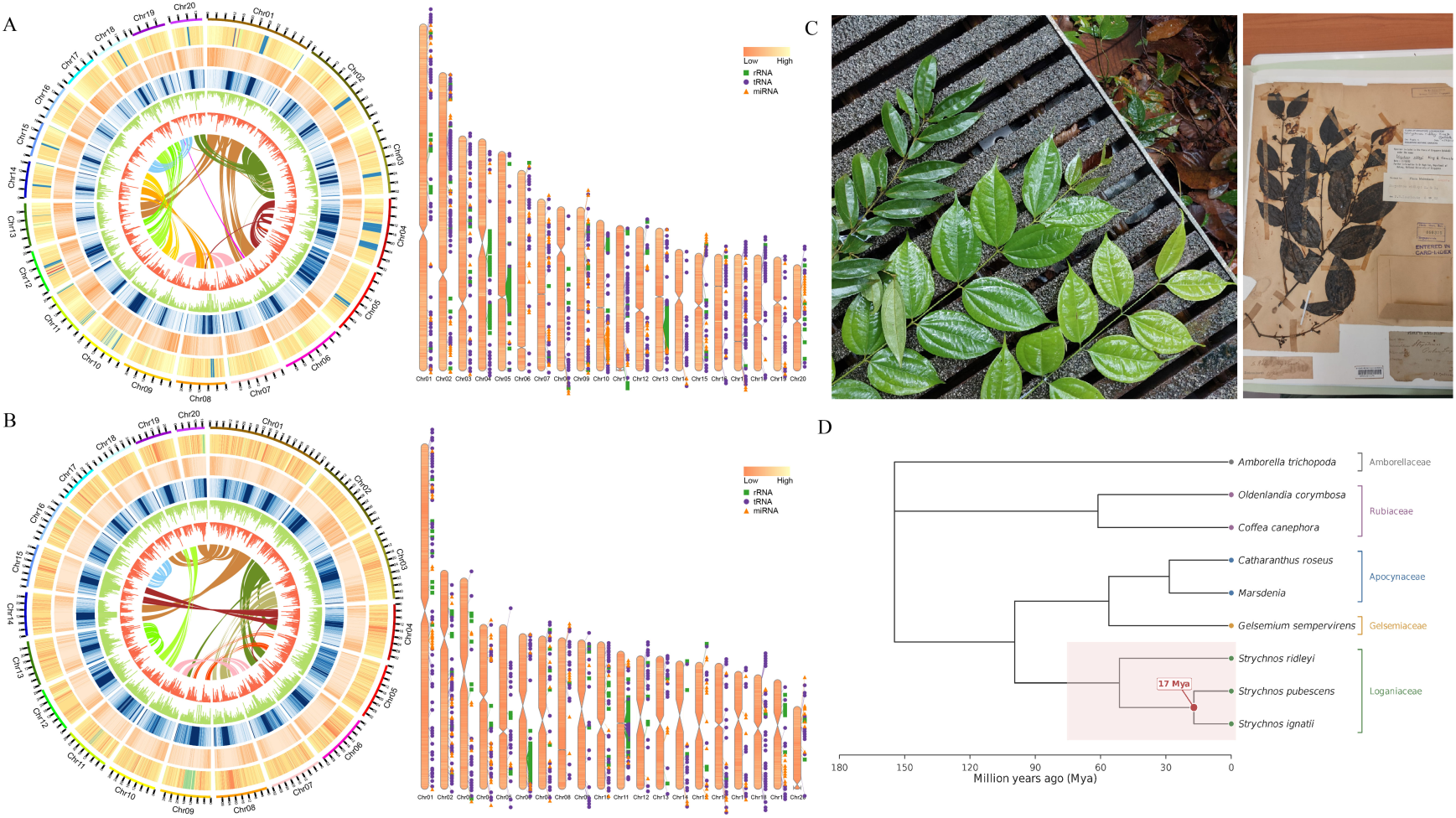
Genome features, phenotypes, and phylogeny of Singaporean Strychnos species. **A–B**: Circos plots showing genome features of S. ignatii (A) and S. pubescens (B). From outermost to innermost, the circles represent GC content, gene counts, repeat content, Copia transposable element (TE) count, and Gypsy TE count, respectively. The lines inside the circles represent syntenic regions between the different chromosomes. Karyoplot illustrates chromosome-wise distributions of genes and non-coding RNAs (rRNA, tRNA, and miRNA). **C**: Left: Leaf comparison of S. pubescens, S. ignatii var., and S. ignatii. Right: A 19th-century herbarium specimen of S. ridleyi collected from the Singapore Botanic Gardens. **D**: Phylogenetic tree reconstructed from 300 shared single-copy orthologous genes across nine species, inferred with IQ-TREE (LG+G4). Strychnos divergence timing was estimated by MCMCTree based on single-copy ortholog alignments identified by OrthoFinder.

To extend these genomic resources to extinct lineages, we reconstructed a draft genome of *S. ridleyi* from a 19th-century herbarium specimen obtained from the Singapore Botanic Gardens (**Fig. 1C**). Illumina sequencing of two preserved leaf fragments yielded approximately 37 Gb of data (∼49× coverage), resulting in a 753 Mb assembly. Despite the degraded nature of the material, the assembly and its 83,820 annotated protein-coding genes achieved relatively high BUSCO completeness scores of 86.7% for the assembly and 84.0% for the annotation, respectively (**Supplementary Fig. S3; Supplementary Table S4**), demonstrating that historical specimens can yield informative draft-quality genome assemblies.

Phylogenetic analysis based on 300 shared single-copy genes from eight Gentianales species and *Amborella trichopoda* placed *S. ridleyi* as sister to the clade comprising *S. ignatii* and *S. pubescens* (**Fig. 1D**), and calibration using *Amborella* fossil data^19^ indicated the divergence of *S. ignatii* and *S. pubescens* at approximately 17 million years ago (**Supplementary Fig. S6**). Alternative allele frequency distributions^20^ revealed a peak at ∼0.33 (**Supplementary Fig. S7**), consistent with elevated ploidy, such as a triploid or hexaploid genome, in *S. ridleyi*. Together, these results establish high-quality genomic resources for *Strychnos*, including an extinct lineage, and provide a foundation for investigating genome evolution and specialized metabolism in Loganiaceae and Gentianales.

### Lineage-specific repeat expansion distinguishes the larger *S. pubescens* genome

Having established the marked difference in genome sizes, we next investigated the underlying structural drivers. Repeat annotation revealed that the smaller *S. ignatii* genome is characterized by a comparatively stable repeat landscape, with repetitive sequences accounting for 43.6% of the assembly (210.5 Mb) and a broadly distributed repeat-age profile rather than a single dominant expansion peak. The predominant repeat classes included long terminal repeat (LTR) retrotransposons (5.88%), particularly Ty1/Copia (3.79%) and Gypsy/DIRS1 (1.64%) elements. The repeat landscape based on Kimura distances displayed a bimodal distribution, with one peak at very low divergence (<2%) consistent with recent but limited transposable element (TE) activity, and a broader older peak at ∼15-25% divergence (**Fig. 2A**).

**Figure 2.**
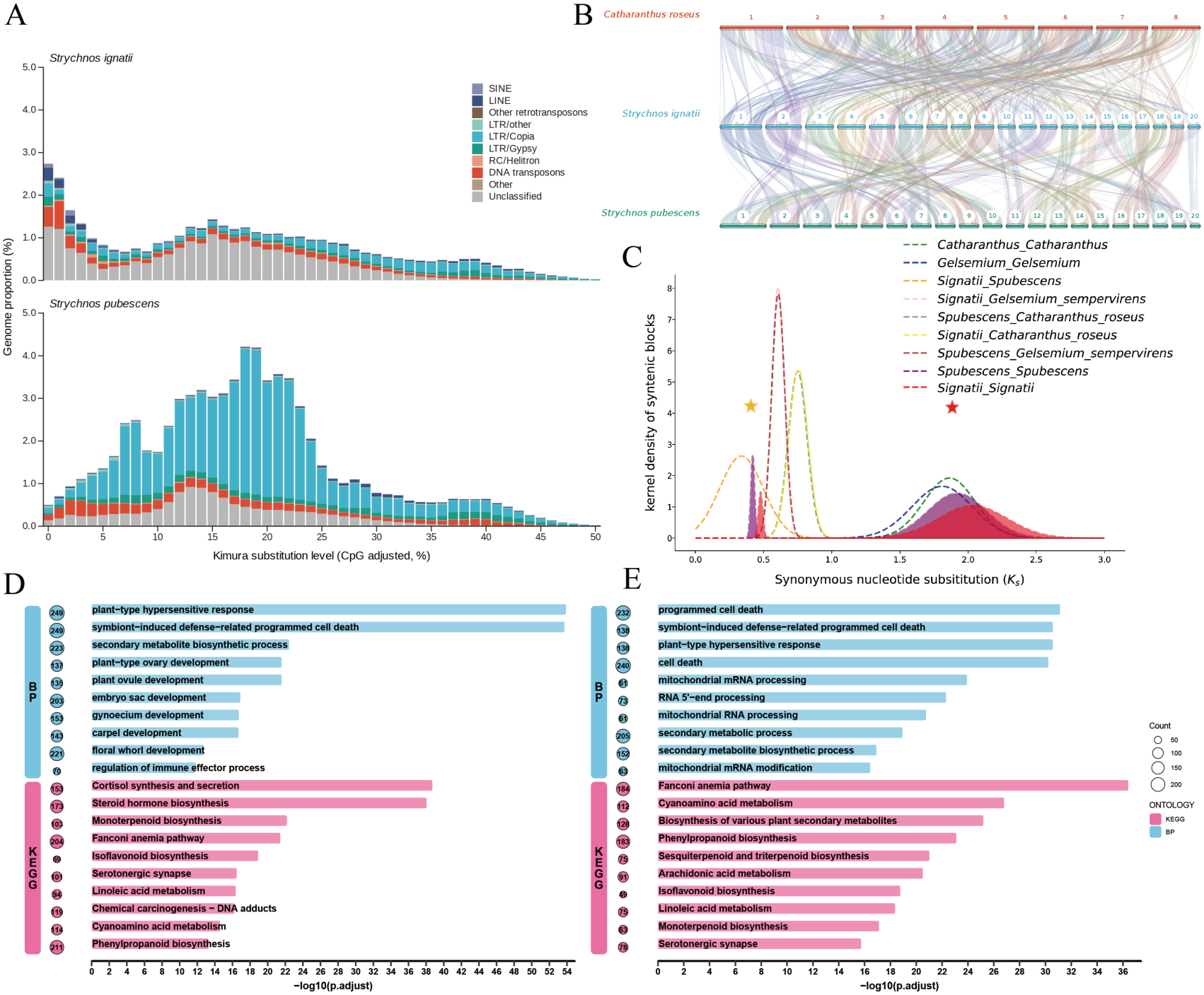
Evolutionary history of Strychnos. **A:** The interspersed repeat landscape and genome fraction of S. ignatii and S. pubescens repetitive elements. **B**: Chromosome-scale synteny among Catharanthus roseus, S. ignatii and S. pubescens. **C**: Density plots of synonymous nucleotide substitution (Ks) values of syntelogs in Strychnos and its closely related species. The red star denotes the gamma paleohexaploid event shared by core eudicots, while the yellow star marks the whole-genome duplication (WGD) event in Strychnos, which occurred approximately 22–26 Mya. **D**: Gene ontology (GO) and KEGG pathway enrichment analysis of tandemly duplicated genes in S. ignatii. **E**: Gene ontology (GO) and KEGG pathway enrichment analysis of tandemly duplicated genes in S. pubescens.

By contrast, the larger *S. pubescens* genome is substantially more repeat-rich, with 54.85% repetitive content. This size discrepancy is driven largely by LTR retrotransposon expansions, as they occupied 22.3% of the assembly, nearly fourfold higher than in *S. ignatii.* The expansion was dominated by Ty1/Copia (17.86%) with a minor amplification of Gypsy/DIRS1 (3.69%) elements. The Kimura divergence profile showed a pronounced unimodal peak centered at 18-20% divergence (**Fig. 2A**). Assuming a neutral substitution rate of 7.77×10^−9^ substitutions per site per generation^21^ and a generation time of five years, this peak corresponds to a major wave of LTR amplification approximately 10-12 million years ago. Because this expansion postdates the inferred divergence of *S. ignatii* and *S. pubescens* (∼17 Mya), this burst represents a major lineage-specific restructuring of the *S. pubescens* genome, potentially generating regulatory sequence variation that could contribute to *cis*-regulatory divergence between the two species (**Fig. 2B**)^22^.

Strikingly, this lineage-specific genome expansion did not correlate with an expanded protein-coding repertoire. Highly complete, multi-evidence gene annotations yielded a total of 39,246 predicted protein-coding genes, alongside predictions of non-coding RNA (ncRNA) elements (**Supplementary Tables S7-S9**) in the smaller *S. ignatii* genome, compared to only 35,408 genes in the larger *S. pubescens* assembly. In both species the completeness of the predicted gene space was high, with BUSCO recovery of 96.1% and 95.3% of conserved orthologs (using the eudicots_odb10 BUSCO database; **Supplementary Fig. S3**), and functional annotation against multiple curated databases assigned putative functions to the majority of predicted genes in both species (**Supplementary Tables S5-S6; Supplementary Figs. S8-S9**). This inverse relationship between genome size and gene count suggests that while TE expansion may have driven the physical enlargement of the *S. pubescens* genome, alternative evolutionary mechanisms – such as differential gene retention, gene disruption due to TE activity, or tandem duplication – have shaped the functional coding space of these two species.

### Whole-genome duplication and lineage-specific genome remodeling underpin metabolic divergence in *Strychnos*

We next investigated the role of whole-genome duplication (WGD) in generating genetic redundancy observed in both species. Analysis of synonymous substitution rates (Ks) among syntelogous gene pairs revealed the ancient gamma paleohexaploid event^23^ in both *Strychnos* species and Gentianales (Ks=2.0-2.2; **Fig. 2C**). Using the same mutation rate and generation time estimates as above (7.77×10^−9^ mutations per nucleotide per generation, five-year generation time), this event was dated to approximately 128-142 Mya, in agreement with previous estimates. Crucially, both *Strychnos* species exhibited a second peak at Ks ≈ 0.4-0.45, corresponding to a WGD time of 22-26 Mya. Because the interspecific divergence occurred at Ks values 0.3-0.35 (corresponding to divergence at 19-22 Mya, which broadly aligns with our phylogenomic estimate of ∼17 Mya), the recent WGD occurred in the common ancestor (**Fig. 2C, Supplementary Figs. S10-S11**) of the sampled *Strychnos* lineages.

Notably, the lineage-specific TE expansion in *S. pubescens* occurred substantially after the inferred WGD and species divergence, suggesting that post-WGD genome evolution proceeded through distinct phases involving initial gene retention and subsequent transposable element proliferation. Additional minor peaks at lower Ks values (<0.2) indicate small-scale segmental duplications occurring approximately 10-13 Mya, broadly coinciding with the period of TE proliferation in *S. pubescens*. The timing of the WGD shortly before species diversification suggests that post-WGD fractionation and differential retention of duplicated genes may have provided the genomic substrate for subsequent divergence in both metabolic gene content and regulatory architecture between the two lineages.

Following a WGD, duplicated genes frequently undergo lineage-specific retention or expansion, providing raw material for metabolic innovation. We therefore examined lineage-specific evolution in terms of tandem expansions, as they have been linked with diversification of pathways that enhance environmental adaptation, including secondary metabolism^24,25^. In *Strychnos*, tandem duplications largely explain why the physically smaller *S. ignatii* genome contains more predicted protein-coding genes than *S. pubescens*, as we identified 13,642 tandemly duplicated genes in *S. ignatii*, while *S. pubescens* tandem duplicates comprised only 9,253 genes, accounting for the discrepancy in their total gene counts.

Tandem duplicates are frequently associated with environmental responses and specialized-metabolism diversification; accordingly, the tandemly duplicated genes in *Strychnos* were significantly enriched for functions related to secondary metabolism. However, the two species exhibited starkly contrasting evolutionary trajectories within the broader terpenoid metabolic network. In *S. ignatii,* lineage-specific tandem duplicates were significantly enriched for monoterpenoid biosynthesis, which supplies upstream precursors for MIA biosynthesis, and contractions for sesquiterpenoid and triterpenoid biosynthesis. Conversely, *S. pubescens* demonstrated the reverse pattern, with duplicates being enriched for sesquiterpenoid and triterpenoid biosynthesis, coupled with the contraction of monoterpenoid biosynthesis processes (**Fig. 2D-E**; **Supplementary Figs. S10-S11; Supplementary Tables S10-S12**). These results were also corroborated through the analysis of gene family expansions in an orthogroup-based analysis (**Supplementary Fig. S6**).

Together, these results indicate that although both species share a common WGD origin, subsequent genome evolution followed markedly different trajectories. In *S. ignatii*, extensive tandem duplication expanded the functional coding space and was preferentially associated with monoterpenoid metabolism. In contrast, *S. pubescens* experienced a major lineage-specific LTR retrotransposon expansion following species divergence, resulting in a larger genome but fewer duplicated genes. These contrasting patterns suggest that metabolic diversification in the two lineages proceeded through different genomic mechanisms, with coding-space expansion predominating in *S. ignatii* and transposable-element-associated regulatory sequence expansion potentially playing a larger role in *S. pubescens* (**Supplementary Figs. S12-S13**).

### Contrasting demographic histories in extant and extinct *Strychnos* lineages

Genome-wide population structure analyses revealed pronounced genetic differentiation between the Singapore populations of *S. ignatii* and *S. pubescens*, with no evidence of recent admixture in either ADMIXTURE or PCA analyses (**Fig. 3A-B; Supplementary Table S13**); these results are consistent with the phylogenomic estimate of Miocene divergence at approximately 17 Mya.

**Figure 3.**
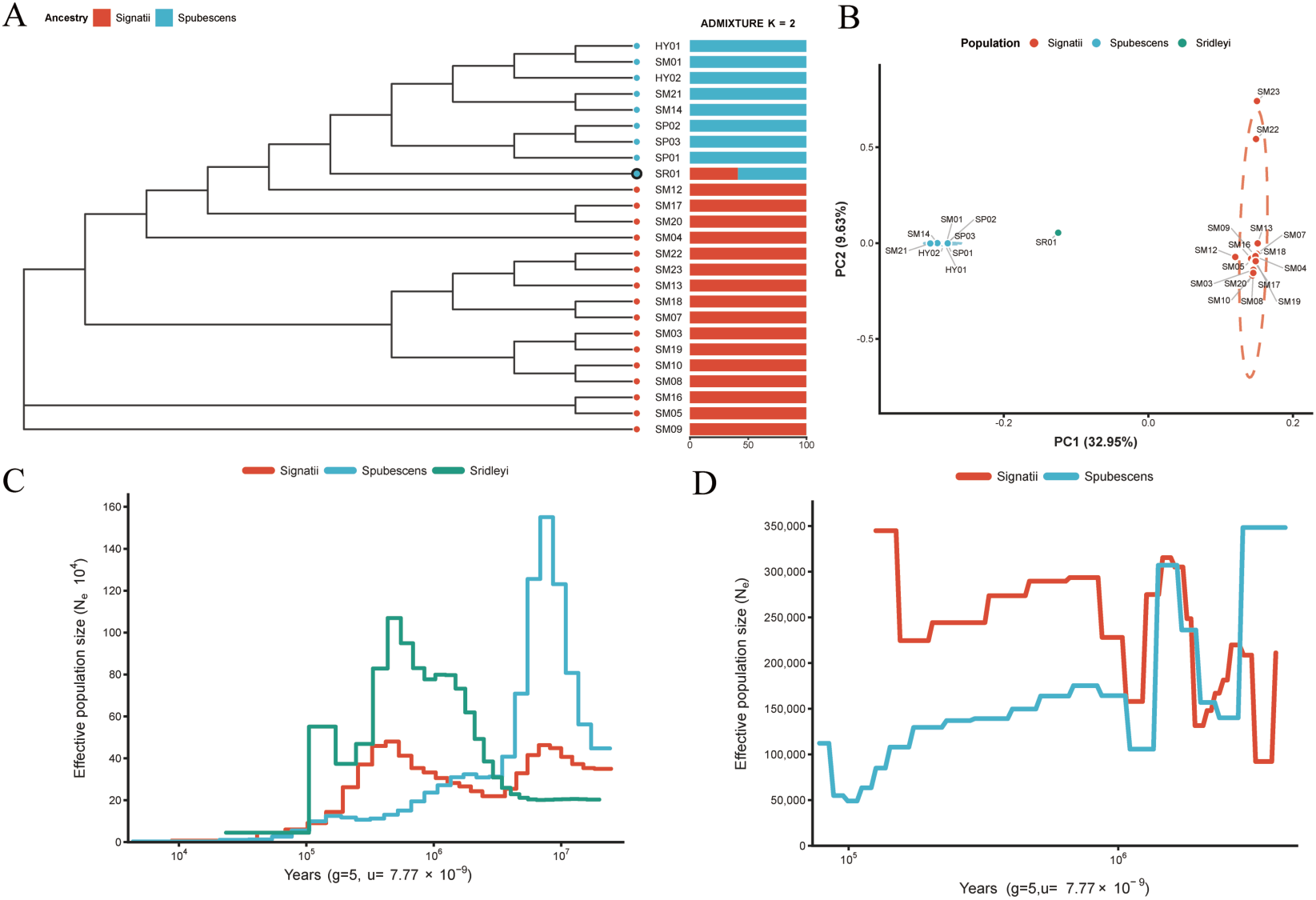
Population genetic analysis of Strychnos. **A**: Genome-wide SNP-based phylogeny and ADMIXTURE analysis of Strychnos samples. K=2 was inferred as the best-supported model, indicating a clear genetic structure between S. ignatii and S. pubescens, with no evident recent admixture between the sampled populations. **B:** PCA analysis of Strychnos based on genome-wide SNPs. The plots indicate that S. ignatii and S. pubescens form distinct clusters along PC1 and PC2, demonstrating clear genetic differentiation between the two species. Additionally, the S. ridleyi individual is positioned between the two clusters. **C**: Historical effective population sizes in the three Strychnos species, estimated using PSMC. **D**: Historical effective population sizes in the two Strychnos species, estimated using SMC++.

Demographic analyses (**Fig. 3C-D**) further indicated contrasting population histories. Historically, *S. pubescens* had a larger inferred effective population size (*N*_e_) than *S. ignatii* around 10 Mya, a period that broadly coincides with the major LTR retrotransposon expansion observed in its genome. However, the inferred demographic trajectories shifted during the Pleistocene, with *S. pubescens* undergoing a marked bottleneck during the Late Pliocene to Early Pleistocene (∼2-3 Mya), followed by a recovery and then repeated bottlenecks from ∼0.8-0.2 Mya. In contrast, *S. ignatii* maintained a larger and more stable effective population size through most of the Pleistocene before declining in the late Pleistocene. Although absolute dates and effective population sizes depend on mutation-rate and generation-time assumptions, these analyses indicate that the two extant species experienced markedly different demographic trajectories after their Miocene divergence, with later population size fluctuations possibly associated with Pleistocene glacial-interglacial climate cycles.

The extinct *S. ridleyi* occupied an intermediate position in both ADMIXTURE and PCA (**Fig. 3A-B)**, consistent with its close relationship to *S. ignatii* and *S. pubescens*. Phylogenetic analyses placed *S. ridleyi* as sister to the clade comprising the two extant species, while its alternative allele-frequency distribution showed a peak at ∼0.33 consistent with elevated ploidy, such as triploid or hexaploid status. PSMC analysis suggested an early effective population size increase, peaking around 10⁵-10⁶ years ago, followed by a prolonged decline without subsequent recovery (**Fig. 3C**). Although the causes of extinction cannot be resolved from genomic data alone, these analyses show that historical herbarium material can recover informative genome-wide signals from extinct plant lineages and provide evolutionary context for comparative studies of *Strychnos*.

### Post-WGD fragmentation and lineage-specific reorganization of the conserved STR–TDC– MATE cluster in *Strychnos*

The *STR*-*TDC*-*MATE* BGC, consisting of *STR*, *TDC*, and multidrug and toxic compound extrusion (*MATE*) genes, is conserved in other MIA-producing Gentianales, including *C. roseus*^28^, *G. sempervirens*^13^ and *O. pumila*^10^. However, syntenic alignments between *Strychnos* and the other known MIA-producers revealed that this BGC is fragmented in both *S. ignatii* and *S. pubescens* (**Fig. 4A-B**). In *S. ignatii*, the *STR* ortholog was found on chromosome 18, whereas the *TDC* ortholog and tandemly duplicated *MATE* paralogs remained colocalised on chromosome 12. In *S. pubescens,* the *STR* ortholog was located on chromosome 12 together with tandemly duplicated *MATE* paralogs, whereas the *TDC* ortholog was colocalised with tandemly duplicated *MATE* paralogs on chromosome 13 (**Fig. 4B**). These syntelogous patterns indicate post-WGD fractionation and lineage-specific reorganization of the ancestral *STR*–*TDC*–*MATE* cluster in *Strychnos*.

**Figure 4.**
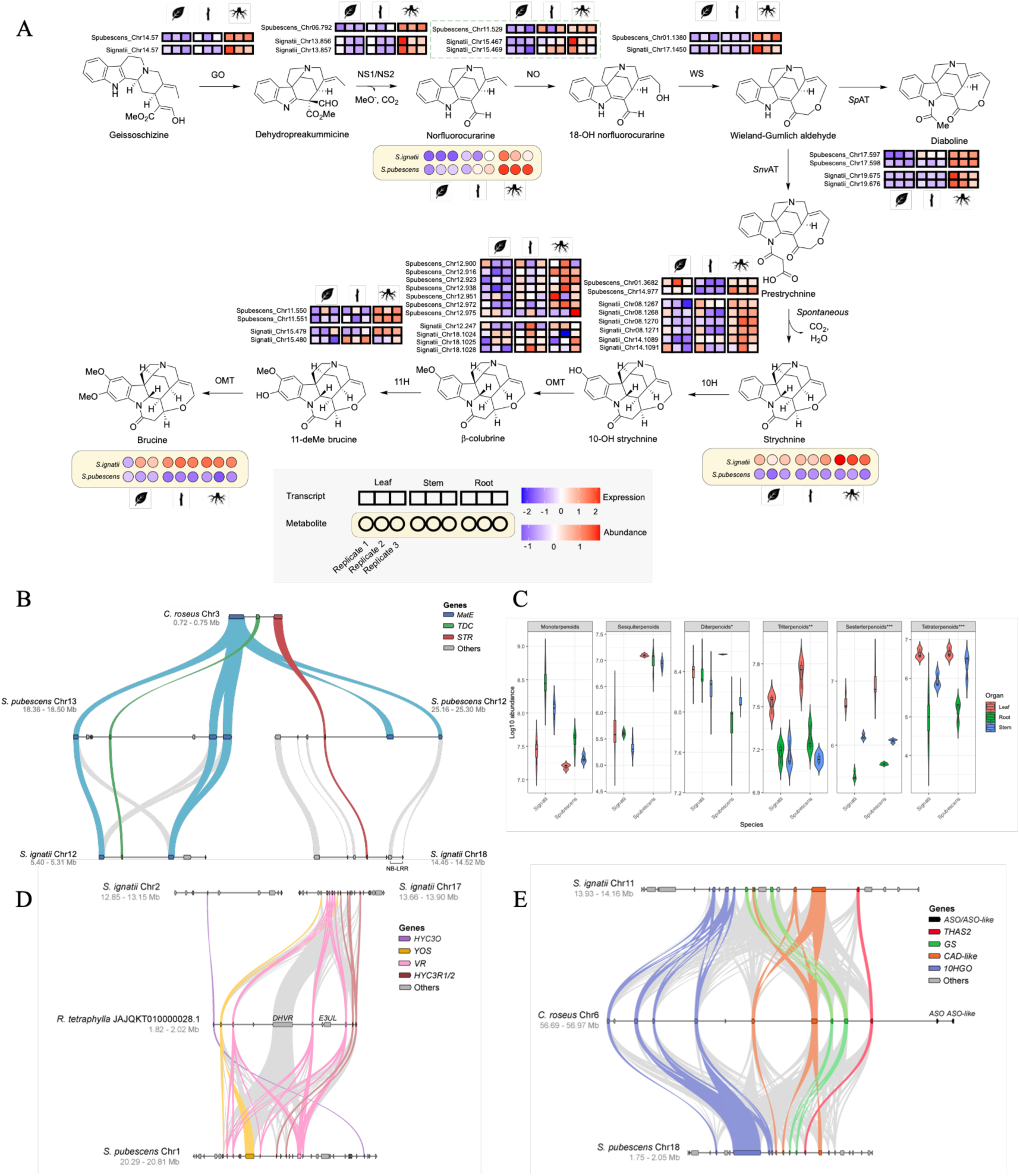
Transcriptomic, metabolomic and syntenic analyses of MIA biosynthesis and biosynthetic gene cluster (BGC) organization in Strychnos. **A**: Strychnine and brucine biosynthetic pathway with heatmaps showing transcript abundance of Strychnos pathway orthologs and metabolite abundance across tissues. Rectangular heatmaps show the standardized TPM (transcripts per million) values for pathway genes in leaf, stem and root replicates of S. ignatii and S. pubescens. Circular heatmaps show metabolite abundance across biological replicates. GO: geissoschizine oxidase; NS: norfluorocurarine synthase; NO: norfluorocurarine oxidase; WS: Wieland-Gumlich aldehyde synthase; AT: acetyltransferase; 10H: strychnine 10-hydroxylase; OMT: hydroxystrychnine O-methyltransferase; 11H: beta-colubrine 11-hydroxylase. **B**: Synteny of the STR–TDC–MATE BGC, showing fragmentation of the cluster across two chromosomes in Strychnos. **C**: Terpenoid-class abundance profiles across root, stem, and leaf tissues of S. ignatii and S. pubescens. Asterisks denote a statistically significant difference between S. ignatii and S. pubescens. One, two and three asterisks indicate P < 0.05, P < 0.01 and P < 0.001, respectively. **D**: Synteny of the reserpine BGC among S. ignatii, R. tetraphylla, and S. pubescens. **E**: Synteny of the geissoschizine synthase BGC among S. ignatii, C. roseus, and S. pubescens.

Despite the fragmentation of the BGC, both species accumulate abundant MIAs, including strychnine, indicating that physical colocalization of *STR* with *TDC* and *MATE* is not required for strictosidine-derived MIA biosynthesis. Transcriptome profiling (**Fig. 4A**) further revealed that although the physical link between the *STR* and *TDC* homologs has been broken in *S. ignatii* and *S. pubescens*, they remain highly expressed in the same tissues, particularly roots. Thus, in *Strychnos*, coordinated expression of the strictosidine pathway has been maintained despite structural disassembly of the ancestral cluster.

### GS and reserpine biosynthetic gene clusters are retained in *Strychnos*

We next asked whether the evolutionary fragmentation observed for the *STR*–*TDC*–*MATE* cluster also extends to other MIA biosynthetic gene clusters. Previous work by Hwang *et al.* identified the geissoschizine synthase (*GS*) cluster as an ancestral feature of Gentianales, whereas the reserpine cluster, containing *HYC3O* and *HYC3R*, was proposed to be a derived feature restricted to the Rauvolfioid clade of Apocynaceae^14,29^. We identified homologous *GS* and reserpine BGCs in the *Strychnos* species (**Fig. 4D-E, Supplementary Figs. S14-S15; Supplementary Tables S14-S15**), extending their distribution to Loganiaceae. Consistent with an ancestral Gentianales origin, the *GS* BGC showed highly conserved organization in both species, comprising homologs of *GS*, *THAS2* and *CAD*-like reductase. The cluster is located on chromosome 11 in *S. ignatii* and chromosome 18 in *S. pubescens* (**Fig. 4E**).

We also identified homologs of reserpine biosynthesis genes *HYC3O*, *HYC3R*, and *VR* in both species. In *S. pubescens*, these genes remain physically linked on chromosome 1, whereas in *S. ignatii, HYC3R* and *VR* are located on chromosome 17 and the oxidase gene *HYC3O* on chromosome 2, indicating lineage-specific fragmentation patterns post-WGD (**Fig. 4D, Supplementary Fig. S15**). Untargeted metabolomic analyses supported reserpine accumulation in both species, particularly in roots, indicating that a reserpine-related biosynthetic pathway is present and active in *Strychnos*. Together, these results show that the genomic architecture and biochemical output of the reserpine pathway are not restricted to rauvolfioid Apocynaceae, but extend to Loganiaceae, where the cluster is retained in *S. pubescens* and fragmented in *S. ignatii*.

To place these findings in a broader genomic context, we performed genome-wide BGC mining using plantiSMASH^30^, identifying 50 candidate BGCs in *S. ignatii* and 53 in *S. pubescens*. Comparative analysis of these predicted clusters identified several conserved and lineage-specific features, including an orthologous terpene-saccharide hybrid candidate cluster (Cluster 39 in *S. ignatii* and Cluster 41 in *S. pubescens*) associated with upstream precursor metabolism. Both genomes also contained alkaloid-related candidate clusters with Berberine Bridge Enzyme-like (*BBE*) domain genes, consistent with their potential roles in oxidative alkaloid chemistry. In addition, *S. pubescens* contained nine BURP-domain-containing cyclopeptide candidate BGCs, several of which span unusually large genomic regions exceeding 1,000 kb, suggesting lineage-specific expansion of peptide-associated specialized-metabolism loci (**Supplementary Tables S16-S17**). However, because these loci remain computational predictions, the strongest evidence for conserved and active MIA-associated BGC evolution in *Strychnos* comes from the *STR–TDC–MATE*, GS and reserpine-related loci described above.

### Comparative metabolomics reveals divergent terpenoid and MIA profiles in two *Strychnos species*

To investigate diversification of specialized metabolism in the two species, we performed untargeted LC–MS profiling across roots, stems and leaves. In total, we detected 530 terpenoid-classified features in *S. ignatii* and 555 in *S. pubescens*, of which 332 were shared between the two species (**Supplementary Fig. S16, Supplementary Tables S18-S19**). Among the shared features, diterpenoids (n=111) and triterpenoids (n=92) were the most abundant classes, followed by terpenoid glycosides (n=41), sesquiterpenoids (n=31), and monoterpenoids (n=27). Across all organs, the overall class composition was dominated by triterpenoids and diterpenoids in both species (**Fig. 4C; Supplementary Tables S20-S31**).

Beyond these shared features, the two species showed marked differences in terpenoid composition. Consistent with gene family expansion analyses, *S. ignatii* displayed a pronounced bias toward higher monoterpenoid accumulation, particularly in roots and stems, whereas *S. pubescens* accumulated higher levels of sesquiterpenoids and triterpenoids (**Fig. 4C; Supplementary Tables S20-S31**). Both species produced strychnine and brucine, with peak abundance in roots, but their accumulation was substantially higher in *S. ignatii*, where root strychnine and brucine levels exceeded those in *S. pubescens* by ∼14,411-fold and ∼1,842-fold, respectively. By contrast, *S. pubescens* was characterized by abundant diterpenoid and triterpenoid features across organs, including oridonin, ursolic acid, hederagenin/soyasapogenol, and lower levels of betulinic acid-, betulin-, and lupeol-type derivatives (**Fig. 4C; Supplementary Tables S20-S31**). These metabolites and related compound classes are widely linked to antimicrobial^31^, anti-herbivory, and insect-deterrent activities^32^, as well as reactive oxygen species (ROS) scavenging and abiotic stress mitigation, suggesting that *S. pubescens* maintains a broad repertoire of specialized metabolites with potential ecological functions.

In addition to strychnine-type MIAs, both species also accumulated other indole alkaloid and MIA-related features, including harmine, serpentine, catharanthine, and vindoline (**Supplementary Fig. S17**). However, these features contributed differently to the overall alkaloid profiles of the two species. In *S. ignatii*, their abundances were significantly lower than strychnine and brucine, whereas in *S. pubescens* harmine accumulated to levels exceeding those of strychnine and brucine (**Supplementary Fig. S18; Supplementary Table S27**).

Together, these results indicate pronounced divergence in the specialized metabolite repertoires of the two *Strychnos* species. *S. ignatii* is dominated by strychnine-centered MIA chemistry, whereas *S. pubescens* exhibits a broader sesquiterpenoid- and triterpenoid-rich profile. These differences are consistent with the contrasting patterns of terpenoid-related gene family expansion identified in the two genomes and provide a biochemical framework for interpreting their distinct metabolic-evolutionary trajectories.

### Comparative pathway analysis identifies norfluorocurarine oxidase as a key divergence point in strychnine biosynthesis

Following the recent elucidation of the complete strychnine and brucine biosynthetic pathway^12^, we identified homologous pathway genes in *S. ignatii* and *S. pubescens* and integrated their expression profiles with metabolite intermediate abundances across tissues (**Fig. 4A; Supplementary Tables S32-S33**). Most pathway homologs were distributed across multiple genomic loci, except for genes residing in MIA-associated BGCs or tandem arrays.

The upstream steps of the pathway were broadly conserved between the two species. The first committed reaction, a four-step oxidation of geissoschizine to dehydropreakummicine catalysed by geissoschizine oxidase (*GO*), was represented by a single ortholog in each genome, with elevated expression in root samples in both species (**Fig. 4A; Supplementary Figs. S19-S20**). The subsequent enzyme, norfluorocurarine synthase (*NS*), was encoded by two tandemly duplicated paralogs in *S. ignatii* whereas *S. pubescens* contained a single homolog. Despite this copy-number difference, expression of both *GO* and *NS* did not differ significantly between the two species, and norfluorocurarine accumulated to comparable levels in roots (**Supplementary Fig. S19**), indicating that precursor supply up to this stage is largely similar in the two lineages.

The major divergence emerged at the step catalysed by norfluorocurarine oxidase (*NO*), a cytochrome P450 that converts norfluorocurarine to 18-OH norfluorocurarine. *S. ignatii* contained two tandemly duplicated *NO* paralogs on chromosome 15, whereas *S. pubescens* possessed a single homolog. Importantly, *NO* expression was significantly higher in the roots, stems, and leaves of *S. ignatii* than in *S. pubescens* (Fold-change = 2.8, P < 0.001; **Fig. 4A**). By contrast, the downstream reaction catalysed by Wieland-Gumlich aldehyde synthase (*WS*) was represented by a single ortholog in each species and showed no significant difference in expression. Together, these results suggest that reduced *NO* expression in *S. pubescens* may limit conversion of norfluorocurarine into 18-OH norfluorocurarine and downstream strychnine pathway intermediates, potentially leaving available precursor to be channelled into alternative metabolic routes (**Supplementary Figs. S19-S20**).

Further downstream, an acetyltransferase (*AT*) converting the Wieland-Gumlich aldehyde into either diaboline or prestrychnine was represented by a single ortholog in both species, with no significant expression differences. In contrast, the late tailoring phase leading to brucine^16^ involved extensive tandem duplication of hydroxylase and methyltransferase genes in both genomes. For instance, *S. ignatii* contained four strychnine 10-hydroxylase (*10H*) tandem duplicates on chromosome 8 and two on chromosome 14, while *S. pubescens* harboured seven *OMT* paralogs colocalised on chromosome 12 (**Fig. 4A; Supplementary Tables S32-S33**). Notably, expression of *10H* was also higher in *S. ignatii* than in *S. pubescens* (fold change = 2.8, P < 0.001). These patterns support a model in which elevated *NO* expression in *S. ignatii* increases pathway capacity toward the strychnine branch, which is then further reinforced by higher *10H* expression. Conversely, low *NO* expression in *S. pubescens* is consistent with reduced pathway capacity toward strychnine and brucine biosynthesis, thereby contributing to the markedly lower accumulation of these alkaloids in this species (**Fig. 4A; Supplementary Figs. S19-S20**).

### *Cis*-regulatory divergence at NO is associated with attenuated strychnine biosynthesis in *S. pubescens*

To investigate the regulatory basis of interspecific variation in strychnine biosynthesis, we performed motif enrichment analyses on the 1-kb promoter regions of all identified pathway homologs. This analysis revealed marked differences in *cis*-regulatory architecture between the two species. The promoter landscape of *S. ignatii* was enriched for motifs associated with transcriptional activation of specialized metabolism, including *de novo* motifs matching *AP2*/*ERF* (MEME-3), *bHLH112*/*OsFBH1* (MEME-1), and *MYB*-related (MEME-2) transcription factors (**Fig. 5A; Supplementary Fig. S21; Supplementary Table S34**). Together, these motifs define a *MYB*, *bHLH*, *AP2*/*ERF* regulatory configuration, consistent with jasmonate-responsive transcriptional programs such as the canonical JA-induced MYC2 (*bHLH*) and ORCA (*AP2*/*ERF*) cascade that regulates MIA biosynthesis in *C. roseus*^33,34^. In plant specialized metabolism, such elements often function as combinatorial jasmonate-responsive modules that integrate stress-related transcriptional inputs^35^. In contrast, the *S. pubescens* promoter landscape was enriched for motifs associated with *TCP* (MEME-6), *BPC* (MEME-1) and *DOF* (MEME-5) transcription factors (**Fig. 5A; Supplementary Table S35**), suggesting a shift toward developmental^36^, chromatin-associated^37^, and abiotic stress-linked^38,39^ regulatory inputs. Consistent with this genome-wide enrichment, DOF-associated motifs were more prevalent across *S. pubescens* MIA pathway promoters than in the corresponding *S. ignatii* promoters, whereas *S. ignatii* retained a broader MYB–bHLH–AP2/ERF-associated promoter architecture across pathway genes (**Supplementary Fig. S21)**.

**Figure 5.**
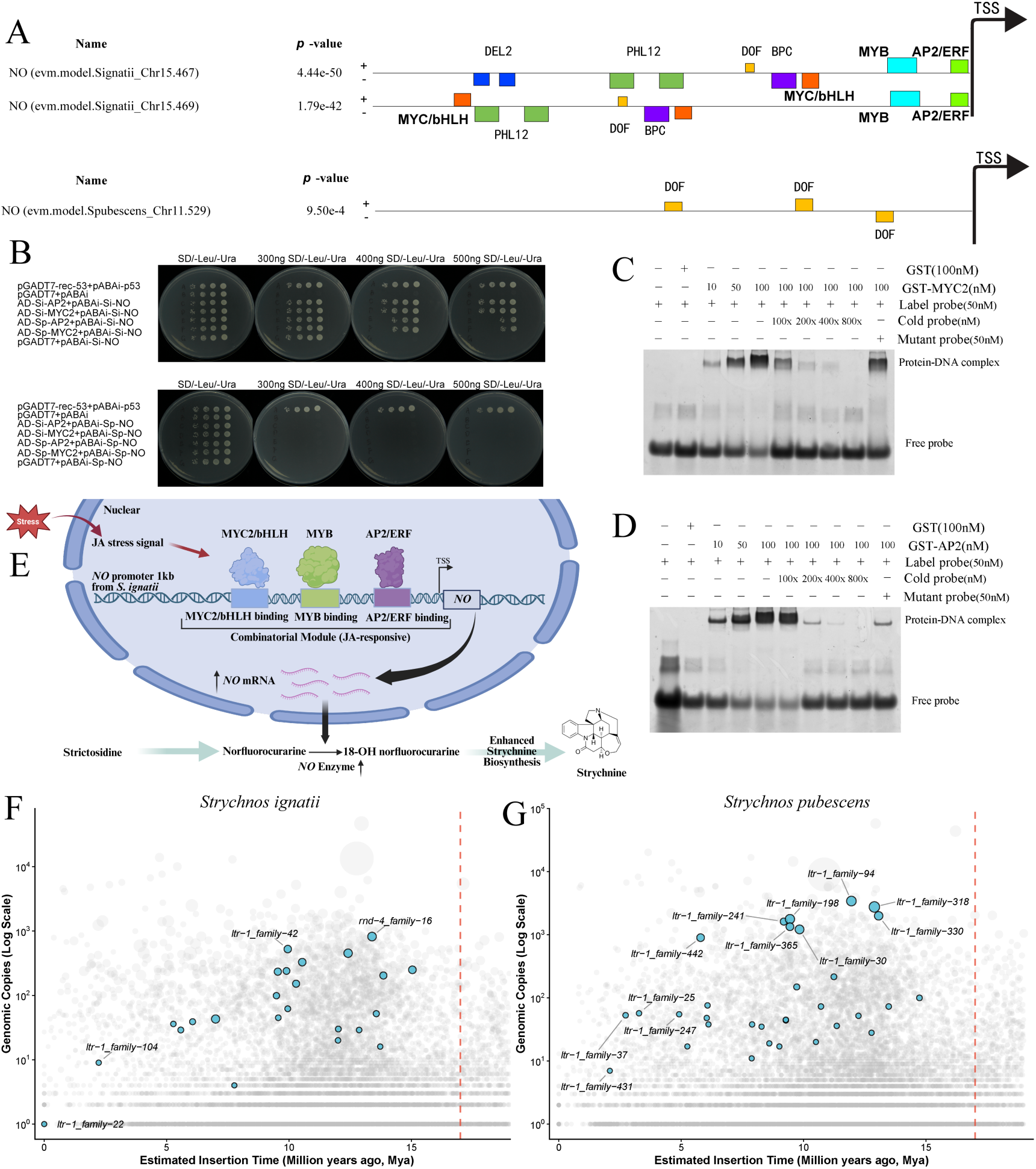
Cis-regulatory divergence at the NO locus supports differential strychnine accumulation between S. ignatii and S. pubescens. **(A)** Schematic representation of significantly enriched transcription factor binding motifs within the 1-kb upstream promoter regions of homologous NO genes in S. ignatii (top) and S. pubescens (bottom). **(B)** Yeast one-hybrid (Y1H) assays testing activation of NO promoter reporters by AP2/ERF and MYC2/bHLH transcription factors. Yeast cells co-transformed with the indicated prey (AD) and bait (pAbAi) vectors were spotted onto SD/-Leu/-Ura media supplemented with varying concentrations of Aureobasidin A (AbA) to assess promoter reporter activation. Empty vectors were used as negative controls. (**C-D**) Electrophoretic mobility shift assays (EMSAs) showing sequence-specific binding of GST-MYC2 and GST-AP2/ERF to the labelled DNA probes containing the corresponding promoter motifs. Each lane contained 50 nM labelled probe, with GST, GST-MYC2 or GST-AP2/ERF protein, unlabelled wild-type (cold) competitor probe, or mutant competitor probe added as indicated above the gel image. Protein-DNA complexes and free probes are marked on the right. **(E)** A proposed working model illustrating the jasmonate (JA)-responsive transcriptional regulation, based on our results and previous work in Catharanthus. In this model, combinatorial MYC2/bHLH-, MYB-, and AP2/ERF-associated regulatory inputs retained in the Si-NO promoter promote elevated NO transcription, contributing to the substantially higher strychnine accumulation observed in S. ignatii. (**F-G**) Amplification dynamics of DOF-associated transposable elements (TEs) in S. ignatii and S. pubescens. Scatter plots showing the estimated insertion time versus genomic copy number (log scale) of TE families in S. ignatii **(F)** and S. pubescens **(G)**. The red dashed line indicates the estimated species divergence time at 17 Mya. Grey circles represent background TEs. Blue circles highlight post-divergence DOF-associated TEs, with circle size proportional to the total genomic sequence occupied by each TE family.

This regulatory divergence was most evident at *NO*, which our pathway analyses identified as a key pathway control point. In *S. ignatii*, the promoters of the two highly expressed *NO* paralogs contain a complex motif repertoire, including *MYC2*/*bHLH* (MEME-1), *AP2*/*ERF* (MEME-3) and *MYB*-binding (MEME-2) motifs, consistent with the canonical JA-inducible transcriptional activation mechanism and elevated pathway expression. By contrast, the single *NO* homolog in *S. pubescens* exhibits a divergent promoter architecture, containing three copies of the DOF-associated (MEME-5) motif but lacking the canonical *MYC2/bHLH*, *AP2*/*ERF* and MYB-associated motifs retained in *S. ignatii* and previously characterized in *Catharanthus* (**Fig. 5A)**.

To experimentally test whether this structural divergence affects promoter activation by candidate transcription factors, we performed yeast one-hybrid (Y1H) assays using the promoter of the highly expressed *S. ignatii NO* paralog (Chr15.467) and its orthologous *S. pubescens NO* promoter (Chr11.529) as bait sequences. We found that *AP2/ERF* and *MYC2/bHLH* proteins from both species activated the *S. ignatii NO* promoter reporter, with the cognate *S. ignatii AP2/ERF* and *MYC2/bHLH* combinations eliciting the strongest growth (**Fig. 5B; Supplementary Table S36**). By contrast, under identical selective conditions none of these four transcription factors were able to activate the *S. pubescens NO* promoter reporter. To further test whether these predicted *cis*-elements are directly recognized *in vitro*, we performed electrophoretic mobility shift assays (EMSAs) using FAM-labeled DNA probes containing the *AP2/ERF* and *MYC2/bHLH* motifs. Probe-only reactions and GST-only controls showed no shifted bands, whereas addition of the target transcription factor proteins produced clear mobility shifts. The shifted signal increased with higher protein input and was progressively reduced by increasing amounts of unlabeled wild-type competitor probes. In contrast, mutated competitor probes failed to abolish the shifted bands, indicating sequence-specific recognition of these promoter motifs (**Fig. 5C-D; Supplementary Table S37**). These Y1H and EMSA assays corroborate our motif-based predictions, demonstrating that the *S. ignatii NO* promoter retains functional *AP2*/*ERF-* and *MYC2/bHLH*-responsive *cis*-elements capable of recruiting pathway-associated activators in heterologous and *in vitro* assays.

Together, our findings support a model in which enhanced strychnine biosynthesis in *S. ignatii* is promoted by the retention of JA-responsive *cis*-regulatory modules at *NO* paralogs, whereas the attenuated strychnine production in *S. pubescens* is associated with the absence of detectable activator-associated *cis*-elements in the analysed *NO* promoter region (**Fig. 5E**). To investigate a possible source of this regulatory divergence, we screened TE families for transcription-factor binding motifs. Several TE families that underwent extensive lineage-specific expansion in *S. pubescens* carried DOF-binding motifs and proliferated during the same 8-13 Mya interval as the major LTR burst, reaching hundreds to thousands of genomic copies (**Fig. 5F-G**). These results indicate that TE expansion substantially increased the abundance of DOF-associated *cis*-regulatory sequence in the *S. pubescens* genome and may have provided a substrate for the emergence of the DOF-dominated promoter architecture observed at the *NO* locus.

### Integrated transcriptome–metabolome analyses recover known MIA genes and identify additional pathway candidates

To identify additional genes and metabolites associated with strychnine biosynthesis, we integrated transcriptomic and untargeted metabolomic data from roots, stems, and leaves of *S. ignatii* and *S. pubescens* using weighted gene coexpression network analysis (WGCNA), regularized canonical correlation analysis (rCCA) and partial least squares (PLS) modelling. WGCNA grouped known MIA-pathway genes into a limited number of coexpression modules, whereas metabolite co-abundance analysis assigned strychnine and related compounds to a small set of metabolite modules **(Supplementary Figs. S23-S27)**.

In *S. ignatii*, strychnine pathway genes were distributed across six coexpression modules (**Supplementary Figs. S23-S28**), with a major module (GEbrown) containing *GO*, *WS*, *AT*, *10H*, and *11H*. Strychnine and numerous upstream and branch-point MIA metabolites were concentrated in a single co-abundance module (MEturquoise) comprising 1,716 metabolite features. We therefore used regularized canonical correlation analysis (rCCA) and partial least squares (PLS) modelling to identify the top 10% of genes most strongly associated with strictosidine, nor-C-fluorocurarine, and strychnine, and extracted the consensus set recovered across models (**Fig. 6; Supplementary Fig. S33; Supplementary Tables S38-S39**). This approach repeatedly recovered validated end-pathway enzymes, including *WS* and *10H*, indicating that the integrative analysis identified biologically relevant pathway-associated gene sets.

**Figure 6.**
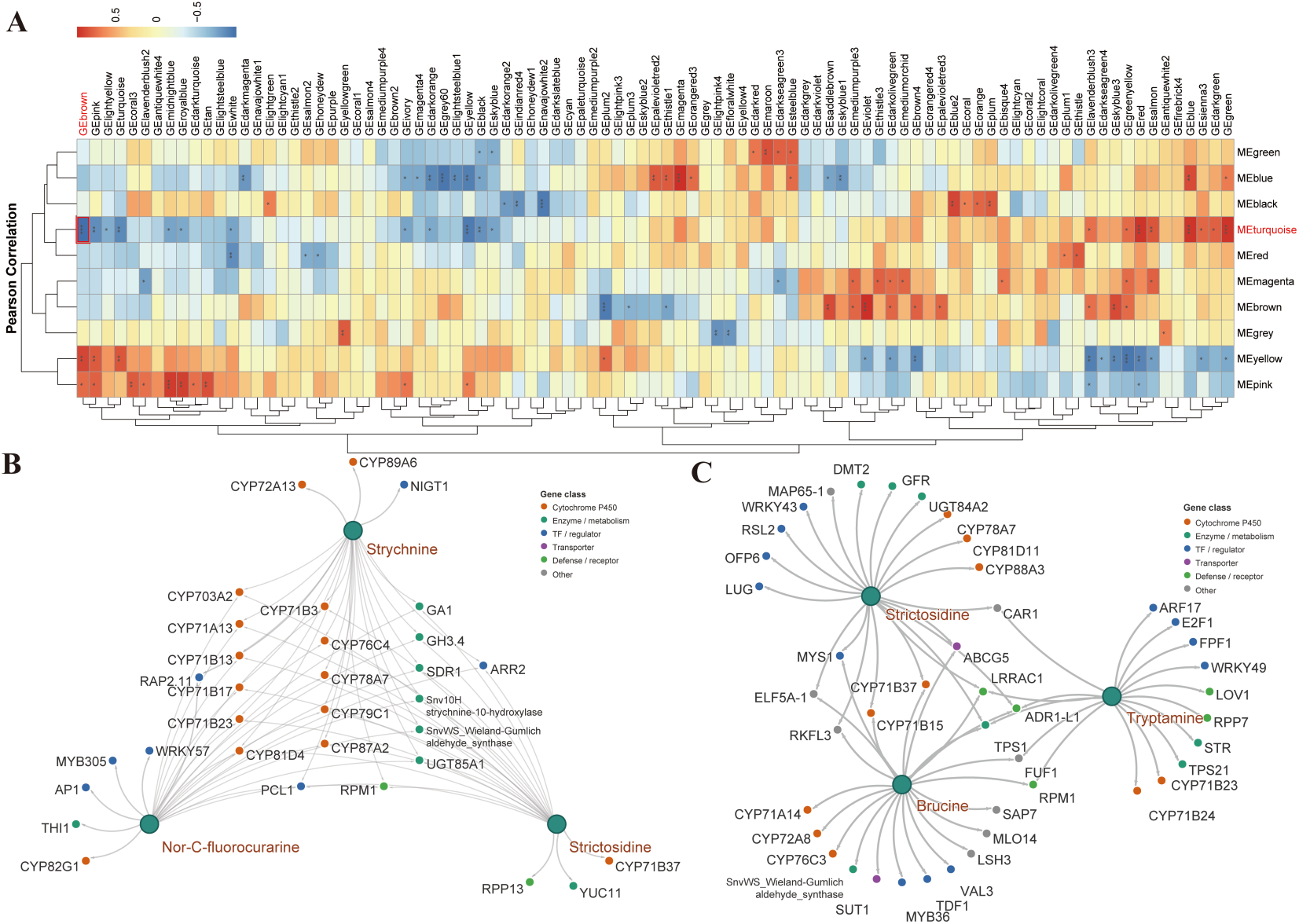
Integrative transcriptome–metabolome analyses identify genes associated with MIA biosynthesis in Strychnos. **A**: Correlation heatmap between metabolite and transcript WGCNA modules. The horizontal axis represents metabolite modules and the vertical axis represents transcript modules. B: Network of candidate MIA-related genes in S. ignatii identified by cross-referencing rCCA and PLS models for the metabolites strychnine, strictosidine, and nor-C-fluorocurarine. Red nodes represent genes identified by both rCCA and PLS, yellow nodes represent rCCA-specific genes, and green nodes represent PLS-specific genes. **C**: Corresponding network of candidate genes in S. pubescens identified using rCCA and PLS analyses for the same three metabolites, with the same color scheme for gene classification.

The *S. ignatii* candidate set was significantly enriched for monoterpenoid biosynthesis and related specialized metabolic pathways, and included enzyme classes associated with oxidative, glycosylation and redox tailoring reactions in MIA metabolism (**Fig. 6B**). These comprised members of the *CYP71* and *CYP76* families^40^, a *CYP82D*-type *10H*^41^, *UGT85A* glycosyltransferases^42^, and short-chain dehydrogenase/reductases (*SDRs*)^43^, all consistent with known chemistry of MIA pathway modification. In addition to catalytic candidates, the intersected sets contained multiple transcriptional regulators, including *WRKY*, *MYB*, and *AP2*/*ERF* family members^44^, as well as stress-associated genes (including uncharacterized *CYP71B* and *CYP71A* clade members), consistent with a link between jasmonate-responsive regulation, defense signalling and MIA output in *S. ignatii* (**Supplementary Tables S40-S42)**.

In *S. pubescens*, core pathway genes were assigned to two main transcript modules, one containing *GO*, *NS*, and *11H*, and the other containing *WS*, *AT*, *STR*, and *TDC*. In metabolite space, the organization resembled that of *S. ignatii*, with tryptamine, strictosidine, and brucine assigned to the same co-abundance module (**Supplementary Figs. S28-S34**). Integrative rCCA and PLS analyses again recovered known pathway components, including *STR* and *WS* homologs, and enrichment analysis identified overrepresentation of plant specialized metabolism pathways. Consistent with *S. ignatii*, multiple members of the *CYP71A*/*B* and *CYP81D* families appeared in the highly correlated gene sets (**Fig. 6C**; **Supplementary Tables S40-S41**), in line with the established role of diverse P450 enzymes in oxidative tailoring of MIA intermediates such as geissoschizine^12,43^.

Across both species, highly associated candidate sets repeatedly included members of the *CYP71*, *CYP72*, *CYP76* and *CYP81* families, together with *UGT* and *SDR* genes, suggesting a conserved enzymatic network for oxidative diversification and downstream tailoring of MIA intermediates^45^. Cross-species comparisons further identified repeatedly recovered candidates, including *CYP78A7* and multiple *P450*-, *UGT*- and *SDR*-family members, suggesting a conserved co-expression framework linking defense, specialized metabolism, and development (**Fig. 6B-C**; **Supplementary Tables S42-S43**).

Promoter analysis further identified *MYB*, *MYC2*/*bHLH*, and *AP2*/*ERF* motif combinations in several *CYP*-family candidates associated with strychnine biosynthesis in *S. ignatii* and *S. pubescens* (**Supplementary Tables S44**-S**45).** Together, these analyses recovered validated pathway genes and identified a broader set of candidate enzymes and regulators associated with strychnine biosynthesis, consistent with a broader association between MIA biosynthesis, jasmonate-responsive regulation and defense-associated metabolism in *Strychnos*.

## Discussion

### Lineage-specific genome remodeling after WGD shaped metabolic divergence in *Strychnos*

Our results reveal how plant specialized metabolic pathways are reshaped by genome evolution after whole-genome duplication. A lineage-specific WGD in *Strychnos* generated duplicated genomic architectures that were subsequently fractionated, rearranged and differentially regulated during diploidization and lineage divergence. Rather than abolishing pathway function, the fragmentation of the ancestral *STR*–*TDC*–*MATE* cluster was accompanied by retention of active MIA biosynthesis, indicating that physical clustering is not an absolute requirement for functional integration of this pathway. This places *Strychnos* as a useful model for understanding how post-duplication genome restructuring can disassemble ancestral biosynthetic gene clusters while preserving coordinated metabolic output.

Post-WGD, the two extant species followed markedly different evolutionary trajectories, defining two contrasting routes to metabolic divergence. In *S. ignatii*, extensive tandem duplication expanded the functional coding space and was preferentially associated with monoterpenoid and MIA metabolism, coinciding with extreme accumulation of strychnine-type alkaloids. In *S. pubescens*, by contrast, genome expansion was driven primarily by LTR retrotransposons rather than protein-coding genes, expanding regulatory sequence space and potentially increasing opportunities for *cis*-regulatory rewiring. These alternative trajectories suggest that closely related plant lineages can diversify specialized metabolism through different genomic mechanisms: expansion of enzymatic coding space in one lineage and expansion of regulatory sequence space in another.

### BGC fragmentation and regulatory coordination after WGD

The distinct behaviour of MIA-associated BGCs illustrates the evolutionary plasticity of plant metabolic architecture after duplication and diploidization. The *STR*–*TDC*–*MATE* cluster, conserved in several MIA-producing Gentianales, has been fragmented in both *S. ignatii* and *S. pubescens*, with *STR* being separated from the *TDC*–*MATE* genes. By contrast, the GS cluster remains a single structurally conserved array in both species, whereas the reserpine-related locus is retained but shows lineage-specific organization, remaining physically linked in *S. pubescens* and fragmented in *S. ignatii*. These contrasting patterns suggest that MIA-associated clusters have followed different evolutionary trajectories after WGD, ranging from structural conservation to fragmentation and reorganization.

Importantly, fragmentation of physical linkage has not abolished pathway activity. Both species retain MIA biosynthesis, and key pathway genes remain co-expressed despite chromosomal separation. This suggests that *trans*-acting regulatory mechanisms can maintain functional integration after structural disassembly of ancestral clusters^46^. More broadly, these findings support a model in which plant BGCs are not static genomic units, but architectures whose physical organization can be reshaped by duplication, fractionation and chromosomal rearrangement while pathway function is maintained through regulatory coordination. This pattern suggests that the evolutionary stability of a metabolic pathway need not depend on the long-term stability of its original cluster architecture. Physical linkage may facilitate pathway assembly and coordinated inheritance, but once regulatory integration is established, pathway function can persist despite erosion of the ancestral cluster.

### TE-associated DOF expansion and regulatory rewiring of strychnine biosynthesis

Following the fragmentation of physical linkage, pathway integration appears to depend increasingly on regulatory coordination between *cis*-elements and *trans*-acting transcription factors. Our pathway analysis identified *NO* as a major expression-level divergence point in strychnine biosynthesis. In *S. ignatii*, duplicated *NO* paralogs retain promoter architectures enriched for *MYC2/bHLH*, *AP2*/*ERF* and *MYB*-associated motifs, consistent with canonical JA-linked MIA activation previously characterized in *Catharanthus*^33,34^. Our yeast one-hybrid and EMSA assays support this regulatory model by showing that *AP2/ERF* and *MYC2/bHLH* transcription factors activate the *S. ignatii NO* promoter, whereas the orthologous *S. pubescens NO* promoter has reduced responsiveness to these activators. Thus, the extreme accumulation of strychnine-type MIAs in *S. ignatii* is associated not only with coding-space expansion, but also with retention of JA-responsive *cis*-regulatory architecture at a key pathway-control point.

By contrast, the reduced strychnine output of *S. pubescens* does not simply represent pathway decay. While the single *S. pubescens NO* homolog lacks the canonical activator-associated motifs, it shows instead a DOF-dominated promoter architecture. Several TE families that expanded during the lineage-specific LTR burst in *S. pubescens* carry DOF-binding motifs, suggesting that TE proliferation increased the abundance of DOF-associated *cis*-regulatory sequence in this genome. Although direct derivation of the *NO* promoter motifs from TE insertions remains to be demonstrated, these results support a model in which TE expansion provided regulatory substrate for rewiring specialized metabolism after WGD. Under this model, *S. ignatii* retained a canonical JA-linked MIA activation program, whereas *S. pubescens* shifted toward a DOF-associated regulatory architecture potentially linked to different environmental-response inputs.

The estimated timing of the *S. pubescens* TE expansion (∼10-12 Mya) coincides with a period of major climatic reorganization across Asia, including the emergence of modern monsoon systems and increasing monsoon seasonality in Southeast Asia^47^. Although our data do not establish a causal connection, environmental instability has frequently been associated with transposable element proliferation and genome restructuring in plants^48,49^. This temporal overlap raises the possibility that changing Miocene climatic regimes may have provided an environmental context for genome diversification. Within the limits of demographic inference, the coinciding demography is also notable, as *S. pubescens* appears to have maintained a high *N*_e_ during the period of TE expansion. Because large *N*_e_ should increase the efficacy of purifying selection against deleterious TE insertions, the LTR burst is unlikely to reflect relaxed selection alone. Instead, it may reflect increased TE activity and/or retention of insertions that were neutral or contributed regulatory variation, including DOF-associated *cis*-regulatory sequence^50^.

These contrasting genome-evolutionary trajectories are mirrored in considerable biochemical divergence. *S. ignatii* is dominated by strychnine-centred MIA chemistry, whereas *S. pubescens* exhibits a broader sesquiterpenoid- and triterpenoid-rich profile. These differences are consistent with the contrasting patterns of gene-family expansion and promoter architecture observed in the two genomes, and suggest that closely related species can reach divergent specialized-metabolism outputs through different combinations of gene retention, tandem duplication, BGC reorganization and *cis*-regulatory rewiring. The fact that strychnine production is not universal within *Strychnos*, as illustrated by *S. potatorum*^16^, further underscores the lability of chemical adaptation strategies within the genus.

### Ecological implications of root-enriched strychnine accumulation

The ecological implications of these divergent biochemical profiles remain hypotheses requiring direct testing. Root-enriched strychnine accumulation and root-biased expression of pathway genes suggest that MIA biosynthesis in *Strychnos* may have important below-ground functions, potentially mediating interactions with herbivores, nematodes, fungi or bacteria^51^. While MIAs are best known for their role in biotic defense, specialized metabolic pathways can also contribute to abiotic stress tolerance, including enhanced reactive oxygen species (ROS) scavenging during salinity, drought or other abiotic fluctuations^52^. Consistent with this broader role, MIAs and their precursors have been reported to respond to severe oxidative stress in related Gentianales species^8,10,13,53^. Thus, the expanded MIA network and retained JA-linked regulatory architecture in *S. ignatii* may support strychnine-centered chemical defense, whereas the DOF-associated transcriptional rewiring in *S. pubescens* raises the possibility that parts of the pathway have been redirected toward developmental or abiotic-stress-associated contexts. DOF transcription factors have been implicated in plant growth, carbon allocation and responses to water limitation, making them plausible mediators of regulatory coupling between specialized metabolism and environmental-response programs^38,39^. However, the relative contribution of strychnine-type MIAs, broader terpenoid repertoires and DOF-associated regulatory rewiring to fitness in natural environments remains unknown. Direct herbivore, pathogen and abiotic-stress assays will be required to determine how these chemical profiles affect ecological performance.

### H**i**storical genome reveals polyploidy and demographic decline in extinct *S. ridleyi*

Together with the two extant genomes, the *S. ridleyi* assembly shows how historical collections can extend comparative genome evolution into extinct branches of plant diversity. Allele-frequency patterns, phylogenetic placement and population-genomic analyses are consistent with a polyploid and possibly hybrid origin. Although the causes of extinction cannot be resolved from genomic data alone, the combination of inferred polyploidy, possible hybrid origin and prolonged demographic decline suggests that extinct herbarium material can provide important context for understanding the evolutionary dynamics of plant lineages no longer available for direct study. Polyploidy can be associated with meiotic instability and reduced fertility in perennial woody plants^26,27^, raising the possibility that genome-level instability contributed to the long-term demographic history of *S. ridleyi*. However, ecological, historical and demographic causes of extinction cannot be resolved from the available genomic data.

In conclusion, our results show that post-WGD metabolic divergence in *Strychnos* was not driven by simple pathway gain or loss, but by the combined effects of BGC fragmentation, differential coding-space expansion and TE-associated *cis*-regulatory rewiring. More broadly, this study demonstrates how genome duplication can decouple physical cluster architecture from pathway function, allowing specialized metabolic pathways to remain active while evolving new coding and regulatory configurations that generate distinct chemical phenotypes among closely related plant lineages.

## Materials and Methods

### Plant material and sequencing

*Strychnos ignatii* and *Strychnos pubescens* were collected in December 2023, from MacRitchie Reserve, Singapore (1.35°N, 103.83°E). Young, healthy leaves, stems, and roots were sampled from each plant, rinsed with PBS buffer, and immediately frozen in liquid nitrogen. To remove secondary metabolites, the samples were treated with sorbitol wash, and high molecular weight (HMW) DNA was extracted using a nuclei isolation method and a modified cetyltrimethylammonium bromide (CTAB) method. DNA quality and quantity were assessed using a NanoDrop 2000 spectrophotometer (Thermo Scientific) and a Qubit 2.0 Fluorometer (Life Technologies). The extracted DNA was used for PacBio Revio HiFi sequencing. For Hi-C genome sequencing, chromatin was extracted using the Dovetail Omni-C kit (21005G) and sequenced on a NovaSeq 6000 platform.

Additionally, three plants of comparable size were selected for RNA extraction from their roots, stems, and leaves using the FastPure Plant Total RNA Isolation Kit (Vazyme Cat. RC401-01). RNA sequencing was performed using Illumina technology, and untargeted metabolomics analysis was conducted using an Agilent G6540B UHD Accurate-Mass Q-TOF LC/MS system.

For population resequencing of Strychnos, DNA was extracted from young leaves using the FastPure Plant DNA Isolation Mini Kit-BOX2 (Vazyme, Cat. DC104-01), and resequencing was performed using Illumina technology.

### Genome Assembly

The chloroplast and mitochondrial genomes of *S. ignatii* and *S. pubescens* were assembled using OATK (v1.0)^54^ with *Arabidopsis thaliana* organelle genomes as references. Reads mapping to organelle genomes were identified and removed from the HiFi sequencing data using Minimap2 (v2.27-r1193)^55^ and samtools (v1.9)^56^. The filtered HiFi data were used to assemble contig-level genomes for both species using Hifiasm (version 0.19.7-r598)^57^. To identify and remove redundant haplotypes caused by heterozygosity, Purge haplotigs (version 1.0.4, parameter: -a 70)^58^ was applied. *S. ridleyi* DNA was obtained from a 19th-century herbarium specimen from the Singapore Botanic Garden, and subjected to Illumina short-read sequencing. After stringent quality control and removal of microbial reads, the genome was assembled using MaSuRCA (v4.1.0)^59^ and then scaffolded using the chromosome-level assemblies of *S. ignatii* and *S. pubescens* as references. Chromosome-level scaffolds were constructed by integrating Omni-C data with contig assemblies using SALSA2 (v2.3)^60^, Juicer (v1.6)^61^, and 3D-DNA (v201008)^62^. Manual refinement and visualization of the scaffold assembly were performed using JuiceBox^63^. Contaminating bacterial and organelle genome sequences (mapping rate > 80%) were identified and removed using BLASTn^64^. Gaps in the genome assembly were filled using TGS-GapCloser (v1.1.1)^65^ with PacBio HiFi reads. The genome was then polished with PacBio data using NextPolish (v1.4.1)^66^ to obtain the final genome assembly. The telomeres and centromeres were predicted and annotated using QuarTeT (v1.2.4)^67^. Genome quality was evaluated using BUSCO (v5.4.2)^68^ with the eudicots ODB10 dataset and Assembly-Stat for genome statistics.

### Genome Annotation

Based on the high-quality genomes obtained, repeat sequences were annotated using RepeatModeler (v2.0)^69^ and RepeatMasker (v4.2.2)^70^. Genome structural annotation for *S. ignatii* and *S. pubescens* was performed through four approaches: de novo prediction, homology-based prediction, transcriptome RNA-seq analysis, and Helixer (v0.3.6)^71^ deep-learning annotation, and combined by the Evidence module (v2.1.0). Coding sequences (CDS) and protein sequences were extracted using gffread (v0.12.9)^72^, and functional annotation was carried out via BLASTP searches against the nr, UniProt, eggNOG, and KEGG databases (E-value ≤ 1e-5, identity > 80%) using DIAMOND (v2.2)^73^. A Venn diagram was used to evaluate annotation coverage across different databases, and the results were integrated to generate the final functional annotation. Ribosomal RNAs (rRNAs) were annotated using barrnap, transfer RNAs (tRNAs) with tRNAscan-SE (v2.0.12)^74^, and other non-coding RNAs were annotated by comparison with the Rfam database.

### Comparative genomic analysis

We conducted comparative genomic analyses of *S. ignatii* and *S. pubescens* with nine representative species of Gentianales, including *Amborella trichopoda*, *Arabidopsis thaliana*, *C. canephora*, *C. roseus*, and *G. sempervirens*. OrthoFinder (v2.4)^75^ was used to classify protein sequences into gene families based on DIAMOND alignments. Single-copy gene sequences were concatenated and used for phylogenetic tree construction with IQ-TREE. Divergence times were estimated with the MCMCTree program, and the resulting tree was visualized using iTOL (Interactive Tree Of Life). Gene family expansions and contractions were identified with CAFE (v5)^76^. GO and KEGG enrichment analyses of expanded and contracted genes were performed using functional annotation results, with an OrgDB database constructed via the AnnotationForge package and enrichment analysis conducted using ClusterProfiler^77^, applying an adjusted p-value threshold of 0.05 for significance. To focus on plant-specific pathways, the KEGG Plant database was retrieved with the R package KEGGREST to filter mapped orthologies. Enrichment results were visualized using ggplot2^78^. Syntenic blocks were identified using MCScanX^79^ and the Python package jcvi^80^. To investigate the evolutionary history of the *Strychnos* genome, synonymous substitution rates (Ks) of collinear gene pairs were calculated with WGDI^81^, and the resulting distributions were visualized as density plots using ggplot2^78^.

Divergence time between *S. ignatii* and *S. pubescens* was estimated using the Ks value and a neutral mutation rate of 7.77 × 10^-9^ mutations per nucleotide per generation^21^, and a generation time of five years. The calculation followed the standard molecular clock equation^82^:

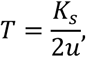

where *T* is the divergence time (in years), *K_s_* represents the peak value of Ks, and *u* denotes the neutral substitution rate.

### Multigene phylogenetic inference and visualization for *S. ridleyi*

We collected protein sequence alignments of 300 complete shared single-copy orthologous genes from 8 Gentianales species, including *S. ignatii*, *S. pubescens*, and S. *ridleyi* from Loganiaceae family, *Oldenlandia corymbosa*, *Coffea canephora* from Rubiaceae family, *Marsdenia tenacissima*, *C. roseus* from Apocynaceae, *G. sempervirens* from Gelsemiaceae and *Amborella trichopoda*. Maximum-likelihood trees were inferred with IQ-TREE under the LG+G4 model. The resulting gene trees were imported into R^83^.

### Strychnos MIA BGCs and biosynthesis genes detection

Based on the known sequences of *G. sempervirens* MIA biosynthetic genes (*STR*, *TDC*, *MATE1*, and *MATE2*)^13^, reserpine BGC (*HYC3O*, *HYC3R*, *YOS, VR*) from *Rauvolfia tetraphylla*, GS BGC from *C. roseus^7^*, and the established strychnine biosynthetic pathway genes (*SnvGO*, *SnvNS1*, *SnvNS2*, *SnvNO*, *SnvWS*, *SnvAT*, *Snv10H*, *SnvOMT*, and *Snv11H*)^12^, BLAST searches were performed against the CDS sequences of *S. ignatii* and *S. pubescens* to identify corresponding homologous genes. These homologs were then located within syntenic blocks, and synteny block analyses of MIA-related genes were visualized with jcvi^79^. BGC homologs detection was performed by CoGeblast and SynFind from CoGe platform (https://genomevolution.org/coge/).

### Resequencing and population genetics analysis

The trimmed reads were aligned separately to the high-quality *S. ignatii* and *S. pubescens* reference genomes using BWA-MEM (v0.7.10-r789). The resulting alignment files were sorted, and duplicate reads were marked with SAMtools (v1.3.1) and PICARD (https://broadinstitute.github.io/picard/). Single nucleotide polymorphisms (SNPs) were then identified using BCFtools^55^. To further refine the SNP dataset, we applied additional filtering with PLINK v1.90b4.6^84^, removing sites with more than 10% missing data and retaining variants with a minor allele frequency (MAF) ≥ 0.01. To reduce the influence of genomic regions exhibiting strong linkage disequilibrium (LD), we conducted a genome-wide LD pruning procedure. Specifically, we scanned the genome using a 50-kb sliding window (advancing every 10 SNPs) and excluded SNPs showing high pairwise correlations (r² > 0.2) within each window.

Population structure analyses were performed using the filtered SNP dataset. Principal component analysis (PCA) was conducted with PLINK v1.90b4.6^84^, and population ancestry was inferred using ADMIXTURE^85^ by testing K values from 1 to 5 and selecting the optimal K based on the lowest cross-validation error. A maximum-likelihood (ML) phylogenetic tree was constructed using RAxML-NG v. 1.0.3 with 1,000 bootstrap replicates to assess branch support. The resulting tree was visualized using iTOL (https://itol.embl.de/), and PCA and ADMIXTURE plots were generated in R^83^.

### Population history and introgression analysis

The pairwise sequentially Markovian coalescent model (PSMC)^86^ was estimated from mapped reads after masking out repeat regions, using standard parameter settings (N25 -t15 -r5 -p 4 + 25*2 + 4 + 6) to estimate the population history. To convert the scaled times and population sizes into real times and sizes, a mutation rate of 7.77 × 10^-9^ mutations per nucleotide per generation^21^ and a generation time of five years were utilized.

Individuals of *S. pubescens* identified through population genetic analysis were mapped to the *S. pubescens* genome to construct a SNP dataset, while individuals of *S. ignatii* were mapped to the *S. ignatii* genome to build a corresponding SNP dataset. SMC++ v1.15.4^87^ was used to infer the demographic histories and divergence times of *S. ignatii* and *S. pubescens* separately. The generation time and mutation rate parameters were set consistently with those used in the PSMC model.

### Strychnine biosynthetic pathway genes regulator analysis by MEME and FIMO

We focused on the promoter regions of strychnine biosynthetic pathway genes in *S. ignatii* and *S. pubescens*, including upstream amine precursor-related genes such as *TDC* and *STR*, as well as downstream candidate Strychnine biosynthesis enzymes. For each gene, we extracted a 1 kb nucleotide sequence upstream of the transcription start site (TSS), retaining strand specificity, and compiled species-specific sequence sets. Motif discovery was performed using the MEME Suite v5.5.8 MEME program^88^. Statistical significance was evaluated with E-values (threshold ≤ 0.05). The top 8 significant motifs (MEME-1-8) from each species were then compared against the JASPAR 2024 CORE plants non-redundant v2 database using Tomtom, with default ungapped alignment and correlation-based distance metrics. Significant matches (q ≤ 0.05) were recorded with their corresponding transcription factor (TF) families/entries.

### RNA-seq data analysis

RNA-seq reads from the leaf, stem and root of three *S. ignatii* and three *S*. *pubescens* biological replicates were aligned to the genomes of *S*. *ignatii* and *S*. *pubescens,* respectively, via STAR (v2.7.10) aligner^89^ using the option quantMode. Gene counts were generated using FeatureCounts. Differential expression analysis was performed on the raw read counts using the R package DESeq2^90^. For the visualization and direct comparison of expression levels, transcripts-per-million (TPM) normalization was applied to the raw counts. Statistical significance of the difference between TPM values for specific candidate genes between two groups was determined using a two-tailed Student’s *t*-test. Expression heatmaps were created using the ggplot2^77^ package of R^83^ and chemical structures were drawn using ChemDraw.

### Untargeted Metabolome data analysis

0.1 g material from each plant tissue sample were first vacuum-dried and then solubilized in a vortex mixture of 1 mL of 100% methanol, acetonitrile, and water. Metabolites were released through physical pulverization and sonication in an ice-water bath. Following sedimentation and centrifugation, the supernatant was concentrated to remove solvents and analyzed using a high-sensitivity mass spectrometer. Data-dependent acquisition was performed with LC-QTOF-MS both in the positive and negative mode.

### Preprocessing of metabolomics data

Metabolites abundance data were standardized and normalized to reduce technical variation and potential batch effects. Missing values were imputed using Probabilistic PCA (PPCA) implemented in the pcaMethods^91^ package in R^83^. Data quality after processing was assessed by calculating pairwise Spearman correlations among samples, and the overall structure of the dataset was examined using PCA. Correlation matrices were visualized as heatmaps generated with the cor and ggplot2^78^ packages, while PCA plots were produced using factoextra.

### Identification of Biosynthetic Gene Clusters (BGCs)

To identify and categorize putative secondary metabolite BGCs, the whole-genome assemblies and corresponding structural annotations of *S. ignatii* and *S. pubescens* were analyzed using plantiSMASH (version 2.0.4)^30^.

### Yeast one-hybrid assays

Yeast one-hybrid assays were performed using the Matchmaker Gold One-Hybrid system. Promoter fragments from *S. ignatii* and *S. pubescens* were cloned into the pAbAi vector and integrated into the genome of the Y1HGold yeast strain after linearization with BbsI. Positive bait-reporter strains were selected on SD/–Ura medium, and basal reporter activity was suppressed by determining the appropriate aureobasidin A (AbA) concentration. Candidate transcription factors, including *AP2/ERF* and *MYC2/bHLH* homologues from *S. ignatii* and *S. pubescens*, were cloned into the pGADT7 activation-domain vector and transformed into the corresponding bait strains using the lithium acetate method (**Supplementary Tables S36**). Transformants were selected on SD/–Leu/– Ura medium and then spotted onto SD/–Leu/–Ura medium supplemented with 300, 400 or 500 ng ml⁻¹ AbA. Yeast growth after incubation at 30 °C was used to assess protein–DNA interactions. pGADT7-rec-53/pAbAi-p53 and empty-vector combinations were used as positive and negative controls, respectively.

### Electrophoretic Mobility Shift Assay

EMSAs were performed using FAM-labeled double-stranded DNA probes containing the predicted AP2/ERF and MYC2/bHLH binding motifs from *S. ignatii* (**Supplementary Tables S37**). Binding reactions were carried out in a 10 μL volume containing 50 nM FAM-labeled probe, purified recombinant protein, and 1× EMSA buffer supplemented with Tris-HCl, DTT, BSA, heparin, glycerol and MgCl2. For competition assays, increasing amounts of unlabeled wild-type competitor probes or mutated probes were added to the reactions. After incubation at 22°C for 15 min, samples were separated on a 6% native polyacrylamide gel in TG buffer under low-temperature conditions. Fluorescent DNA–protein complexes were visualized using a Typhoon fluorescence scanner. GST protein and probe-only reactions were included as negative controls.

### Identification and Evolutionary Analysis of DOF associated TEs Regulatory Elements

To investigate the potential transposable element (TE) mediated dispersion of DOF transcription factor binding sites, LTR/Gypsy and LTR/Copia consensus sequences generated by RepeatModeler were screened for the DOF core motif. The insertion time of each TE family was estimated using the same equation 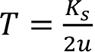, where u represents the synonymous mutation rate of 7.77 × 10^-9^ mutations per nucleotide per generation, and a generation time of five years. An evolutionary landscape of TE insertions was subsequently constructed, mapping the estimated insertion time against the log-transformed genomic copy number, with bubble sizes scaled to the total genomic footprint of each family. The lineage-specific speciation node (∼17 Mya) was incorporated as a chronological reference to delineate recent, post-speciation TE amplification bursts.

### Weighted Gene Co-expression Network Analysis (WGCNA)

Co-expression networks for both transcriptomic and metabolomic datasets were constructed using the WGCNA package^92^ in R^83^. Expression and metabolite abundance matrices were log₂(x + 1) transformed prior to network construction. For each dataset, the soft-thresholding power (β) was selected based on the scale-free topology criterion (R² > 0.8). Adjacency matrices were computed and converted to topological overlap matrices (TOMs), which were used to perform hierarchical clustering. Modules of co-expressed genes or metabolites were defined using the dynamic tree cut method. Network structures within individual modules were visualized using visNetwork and Cytoscape (v3.10.3)^93^.

### Integrated Analysis of Transcriptomics and Metabolomics Data

Module eigengenes derived from transcriptomic and metabolomic WGCNA analyses were used to examine cross-layer relationships. Pearson correlation coefficients were calculated to identify associations between gene and metabolite modules, and the resulting correlation matrix was visualized with pheatmap. To further explore coordinated variation across the two datasets, regularized canonical correlation analysis (rCCA) and partial least-squares (PLS) analysis were performed using the mixOmics package^94^. The shrinkage-based approach was applied to determine optimal lambda parameters for rCCA. Overlapping genes and metabolites identified from these integrative analyses were visualized using Cytoscape (v3.10.3)^93^.

## Data availability

Strychnos PacBio HiFi and Hi-C raw sequence data reported in this paper have been deposited under NCBI Bioproject PRJNA1492247, the transcriptome data have been deposited under PRJNA1496554, and the Strychnos population resequencing raw data have been deposited under PRJNA1492260. Strychnos genome and annotation are shared in figshare: https://doi.org/10.6084/m9.figshare.33252795. Other data supporting the findings of this work are available within the paper and its Supplementary Information files. All data are available from the corresponding author upon reasonable request.

## Author contributions

J.L., B.T., and J.S. conceived and designed the experiments. J.L., J.J., R.A., R.T., and M.N. collected samples. A.L. and W.L. extracted the HMW DNA samples. J.L. performed the experiments. J.L., J.J. and J.S. performed the analysis. J.L., J.J. and J.S. wrote the original manuscript, completed with reviews, edits and revisions by R.A., H.Z., M.N., J.J.N, R.T.J.K, G.K., and B.T. All authors read and approved final manuscript.

## Acknowledgments

Thanks to the High Performance Computing Centre (HPCC) at Nanyang Technological University for providing server administration. Thanks to the Singapore Botanic Gardens and the National Parks for granting permission to collect samples and specimens. This work was supported by the Verdant Foundation, Ngee Ann Kongsi, NCCS Cancer Fund, and Tan Yew Oo Distinguished Professorship, SingHealth Duke-NUS Academic Medical Centre grant: AMRI/BD-MED/FY2024/EX/60-A121, and the Singapore MoE tier 1 (RG82/21), Tier 2 grant (MOE-T2EP30223-0021) to J.S, Academy of Finland (decisions 318288 and 319947 to J.S), and CoE in Tree Biology (TreeBio AoF CoE 346139; 346141).

