## Supplementary Figures for "Whole-genome duplication drives biosynthetic gene cluster fragmentation and regulatory rewiring of monoterpene indole alkaloid metabolism in *Strychnos*"

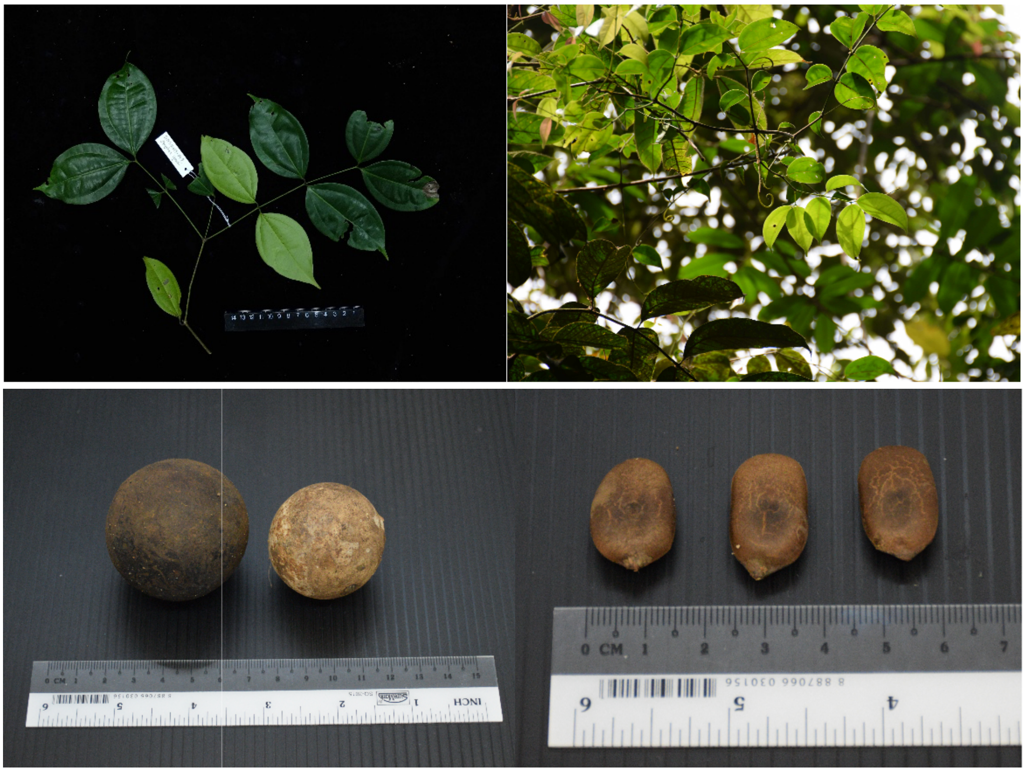


**Figure S1.** Representative morphological traits of *Strychnos ignatii*, including leaves, fruits, and seeds from field collections.

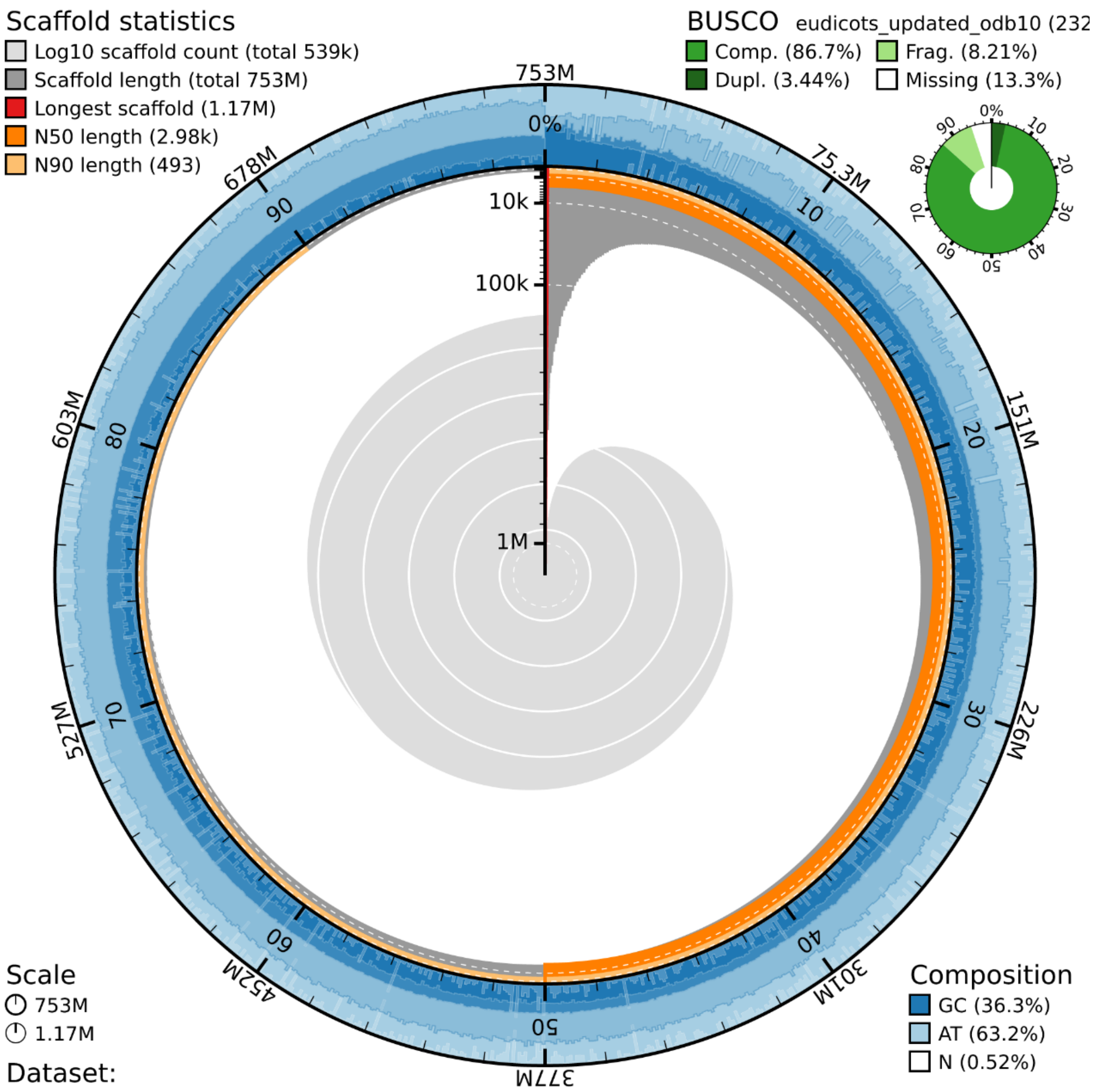


**Figure S2.** Snail plot of *S. ridleyi* genome, illustrating the assembly completeness.

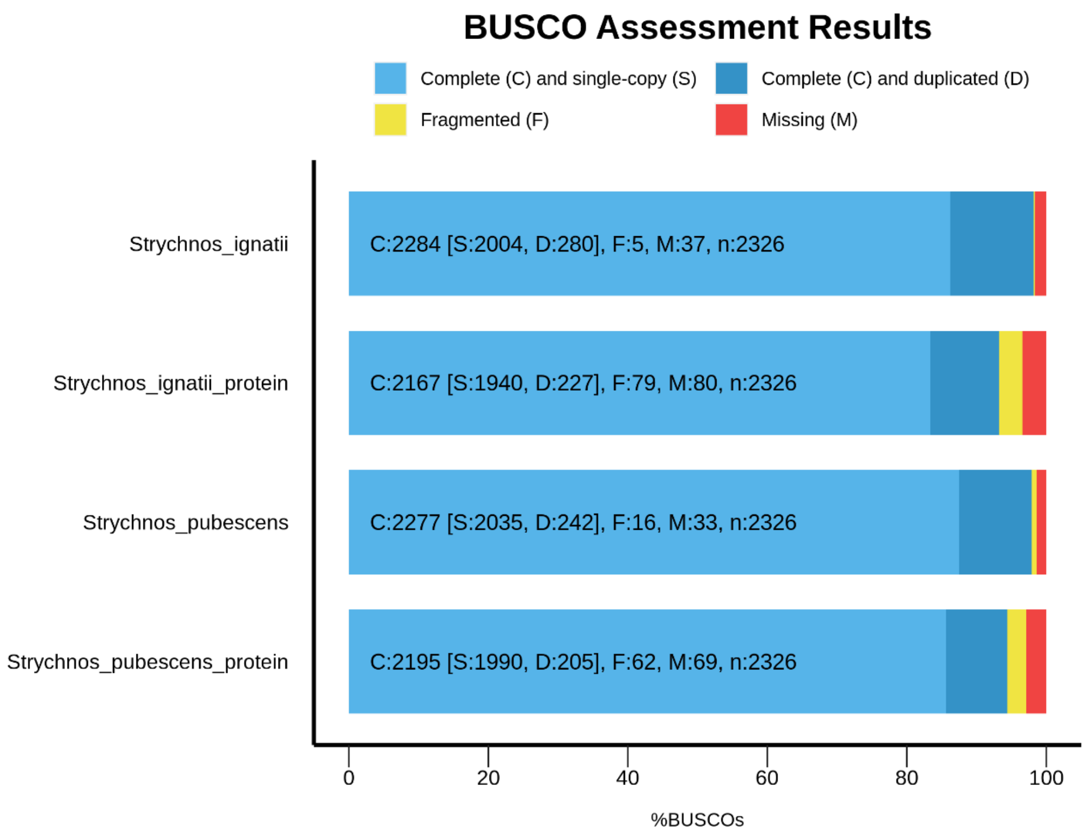


**Figure S3.** Conserved single-copy gene-based (BUSCO) assessment of genome assembly and annotation completeness for the two Strychnos species.

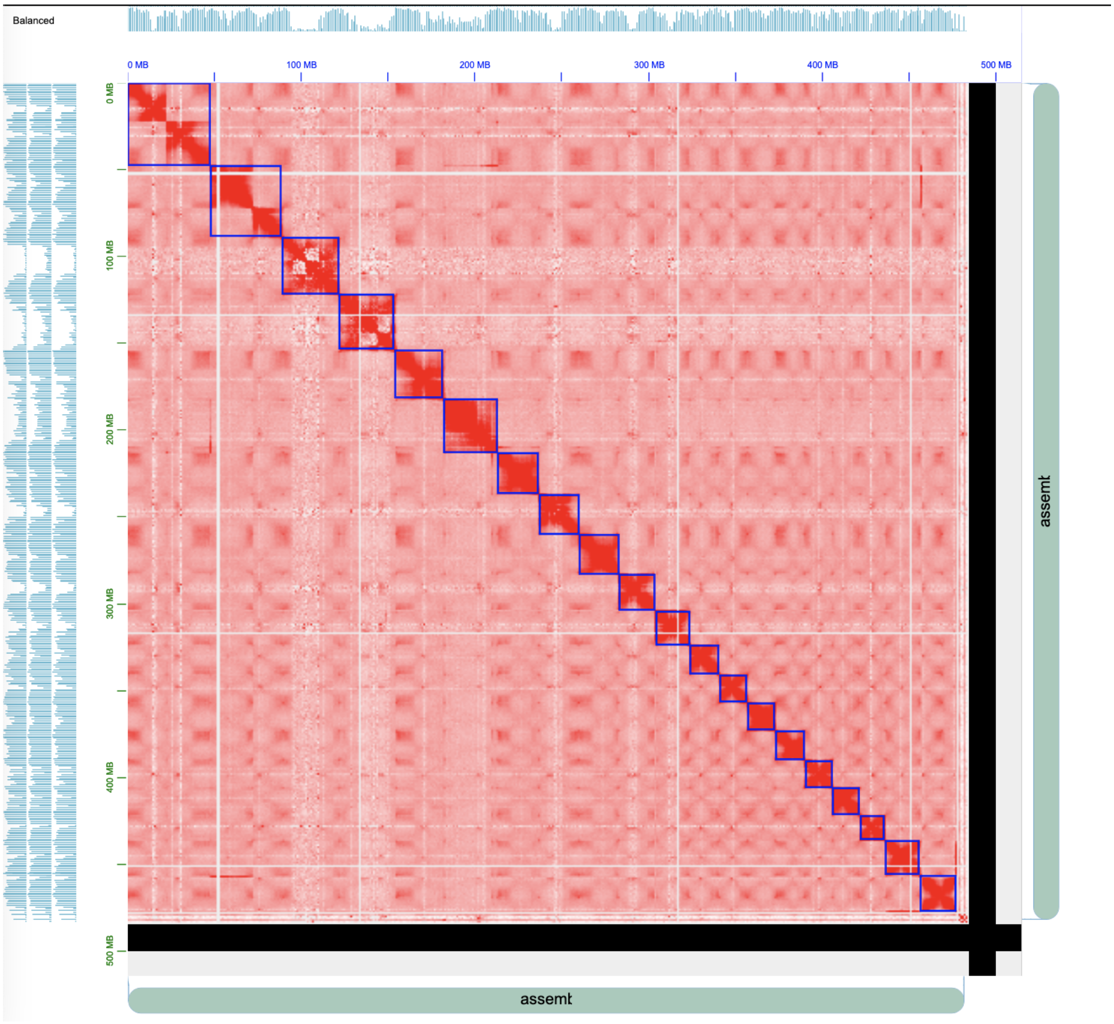


**Figure S4.** The contact map of *S. ignatii* hic assembly by Juicebox.

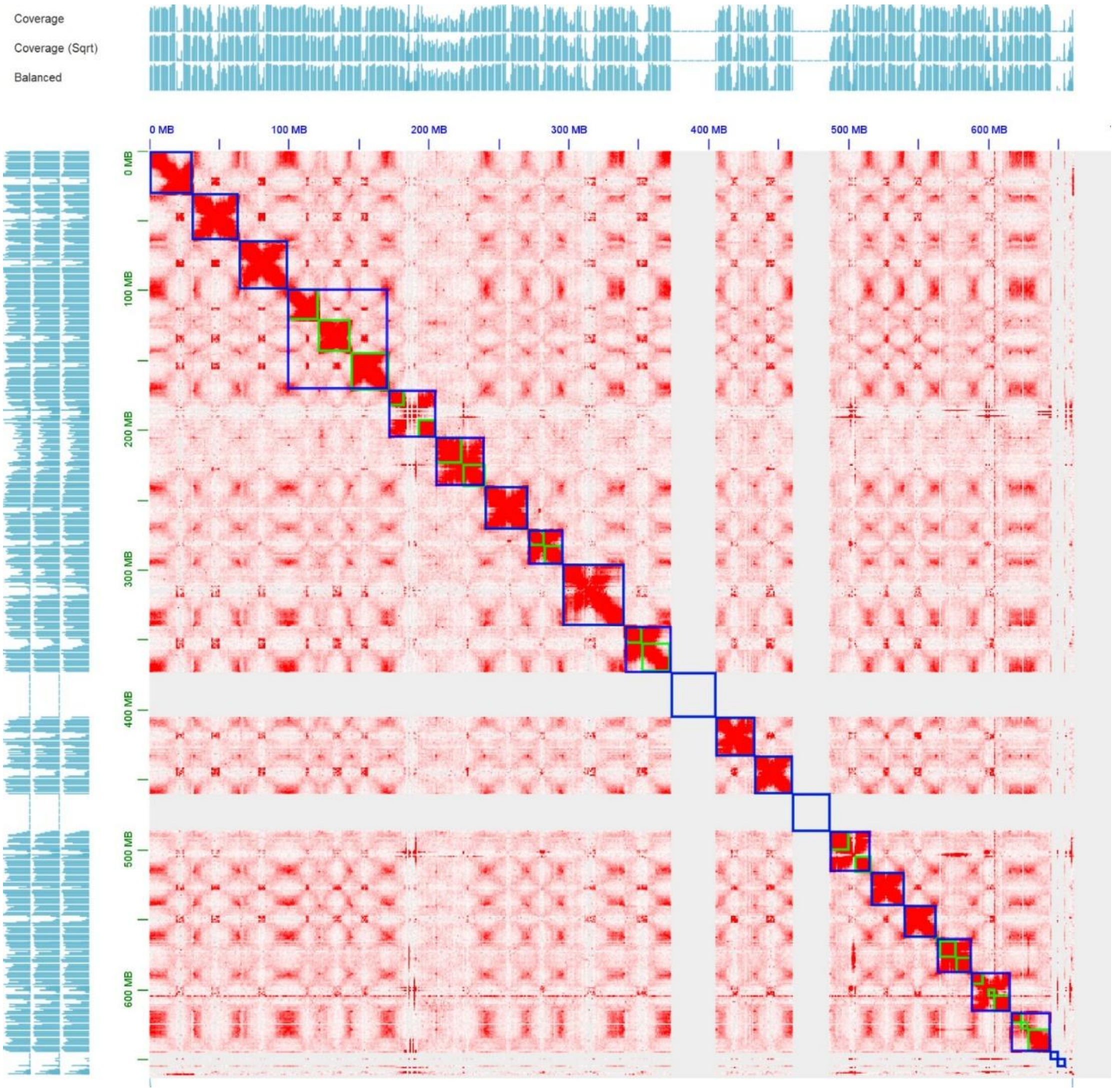


**Figure S5.** The contact map of *S. pubescens* hic assembly by Juicebox.

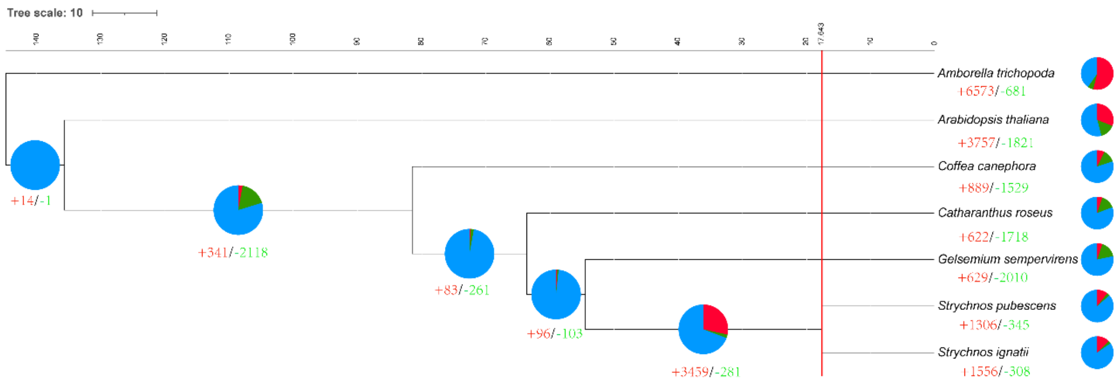


**Figure S6.** Phylogenetic tree of Strychnos within Gentianales based on orthologous groups identified by OrthoFinder, the x-axis shows divergence time scale. The red line marks the divergence of Strychnos at ~17.643 Mya. Pie charts indicate conserved (blue), expanded (red), and contracted (green) orthogroups.

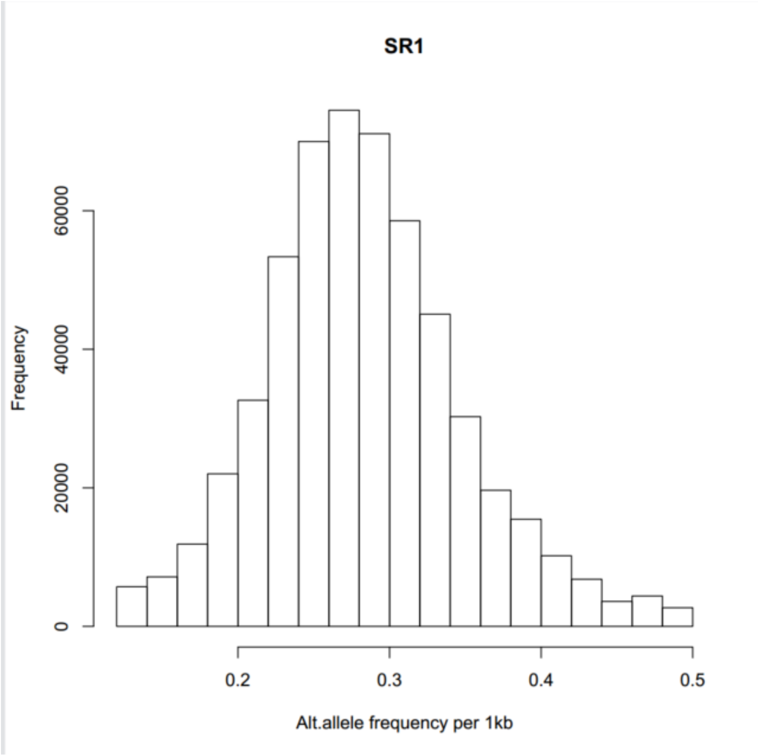


**Figure S7.** Density plot of alternative allele frequency *Strychnos ridleyi*.

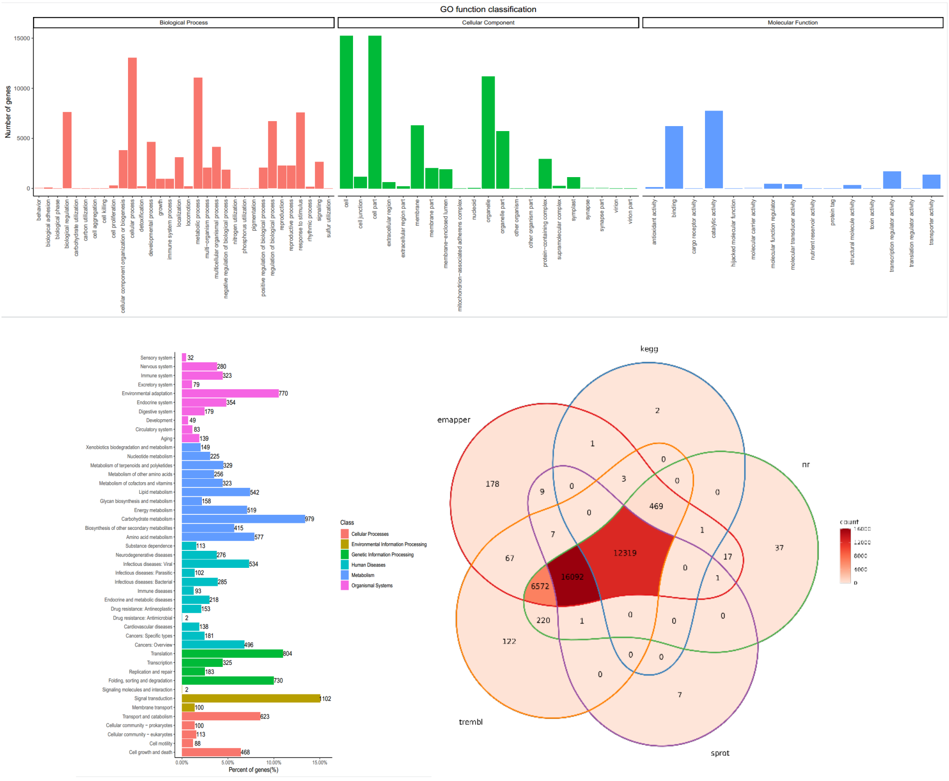


**Figure S8.** Functional annotation results of the *S. ignatii* genome.

Note: The upper figure show the *S. ignatii* GO enrichment analysis, the below figures show the KEGG enrichment results, and the Venn diagram summarizing the functional annotations assigned by NCBI non-redundant (nr), UniProt, eggNOG, and KEGG.

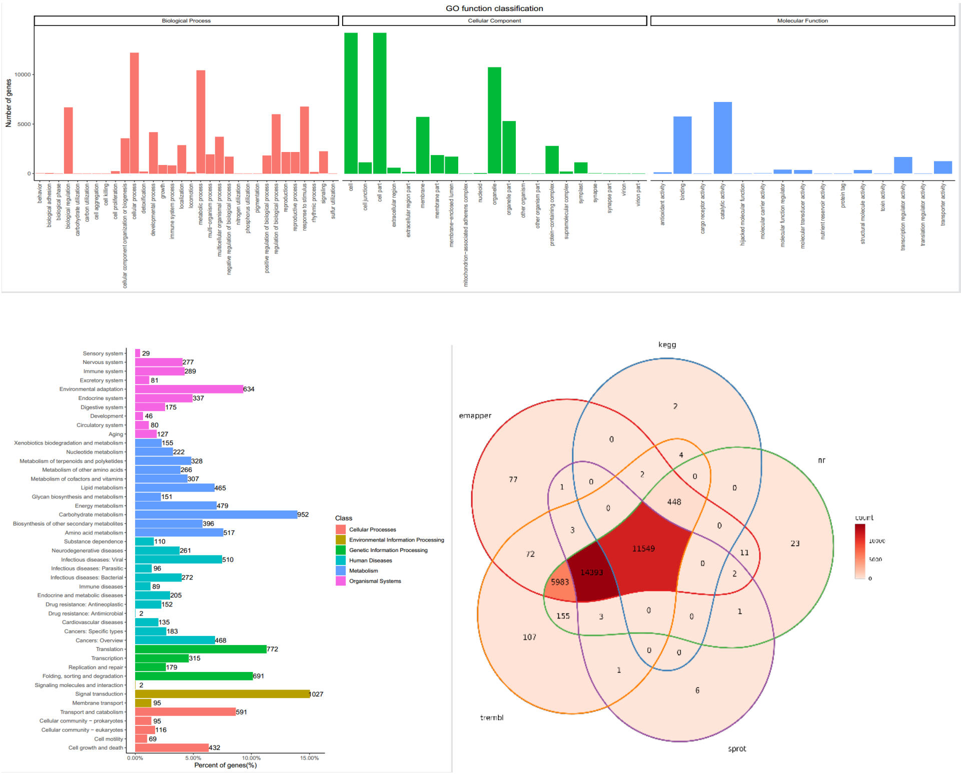


**Figure S9.** Functional annotation results of the *S. pubescens* genome.

Note: The upper figure show the *S. pubescens* GO enrichment analysis, the below figures show the KEGG enrichment results, and the Venn diagram summarizing the functional annotations assigned by NCBI non-redundant (nr), UniProt, eggNOG, and KEGG.

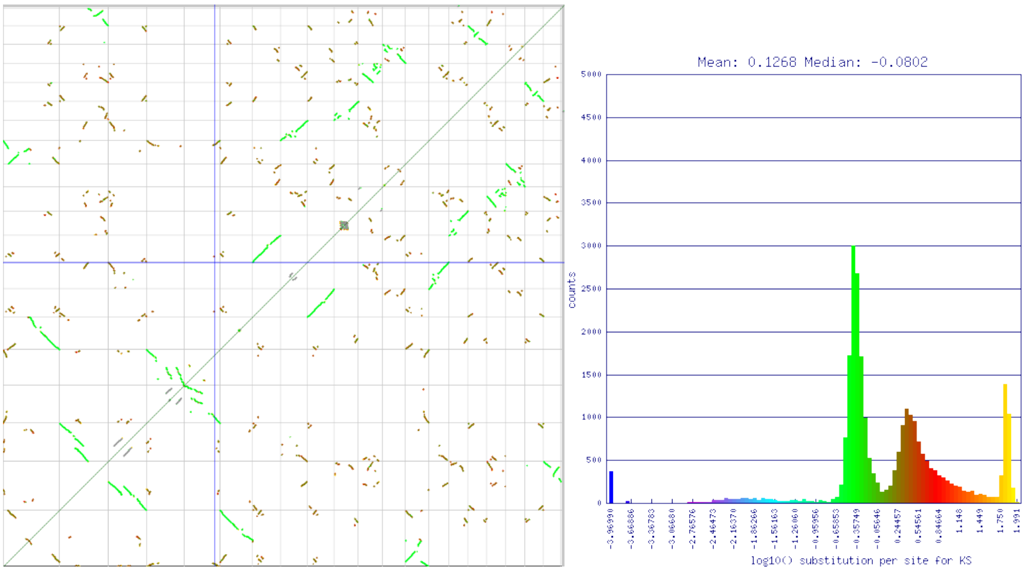


**Figure S10.** CoGe results of the *S. ignatii* self-alignment.

Note: The left panel shows the self-alignment of *S. ignatii* genome, and the right panel displays the histogram of Ks values. The figure below shows the enrichment analysis results of tandem repeat genes.

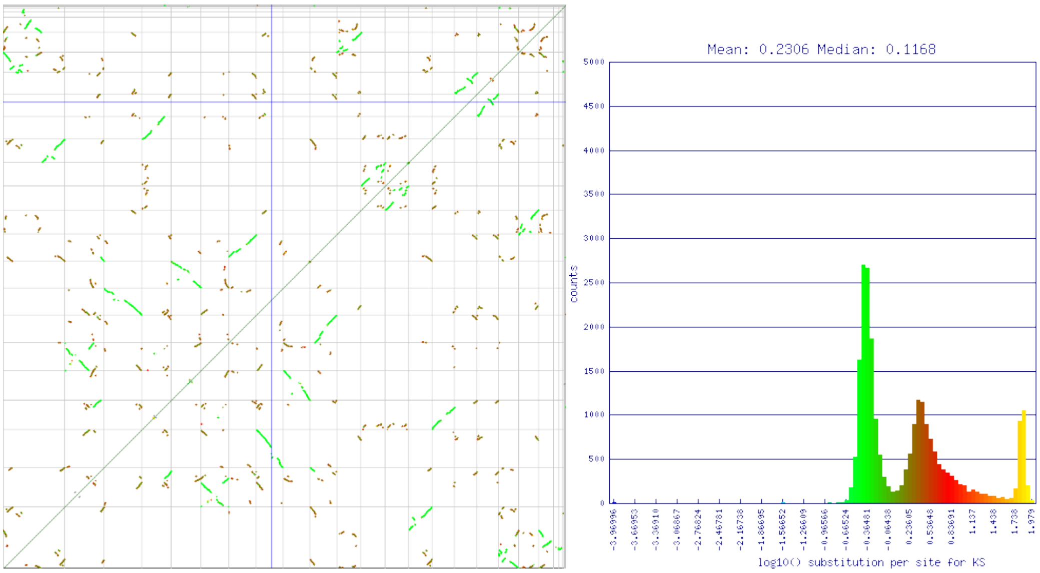


**Figure S11.** CoGe results of the *S. pubescens* self-alignment.
Note: The left panel shows the self-alignment of *S. pubescens* genome, and the right panel displays the histogram of Ks values. The figure below shows the enrichment analysis results of tandem repeat genes.

Significantly enriched gene ontology (GO) and KEGG pathways among significantly expanded orthogroups in S. ignatii. E: Significantly enriched GO and KEGG pathways among significantly expanded orthogroups in S. pubescens.

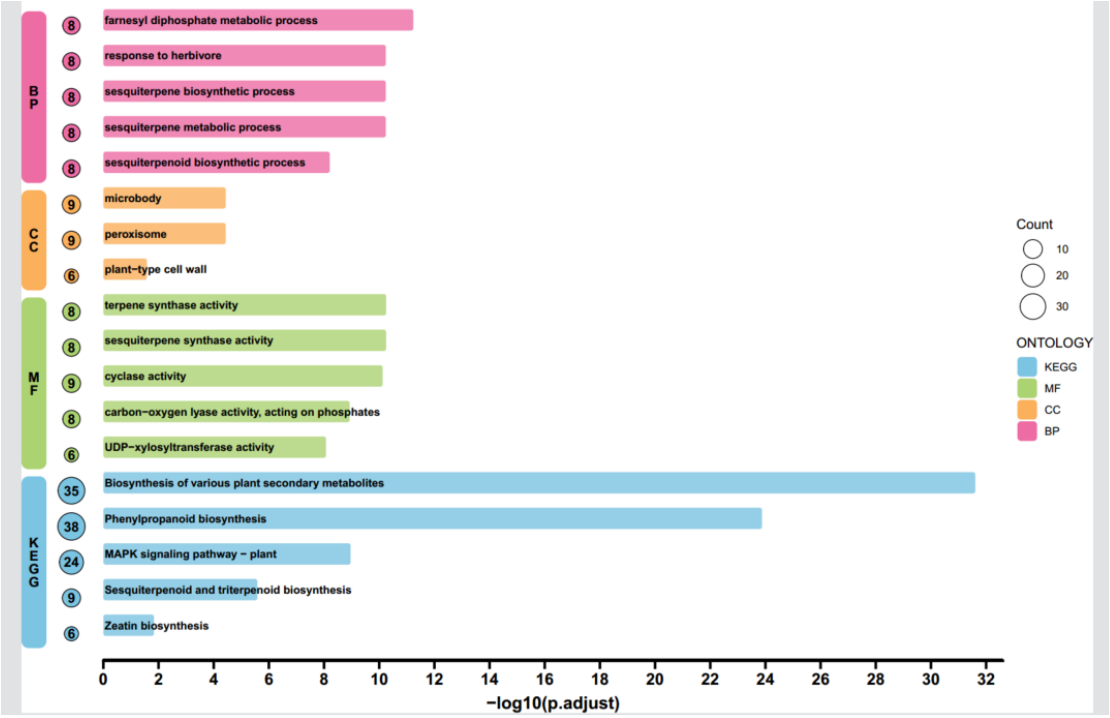


**Figure S12.** Enrichment analysis of *S. ignatii* contracted genes based on the CAFE results.

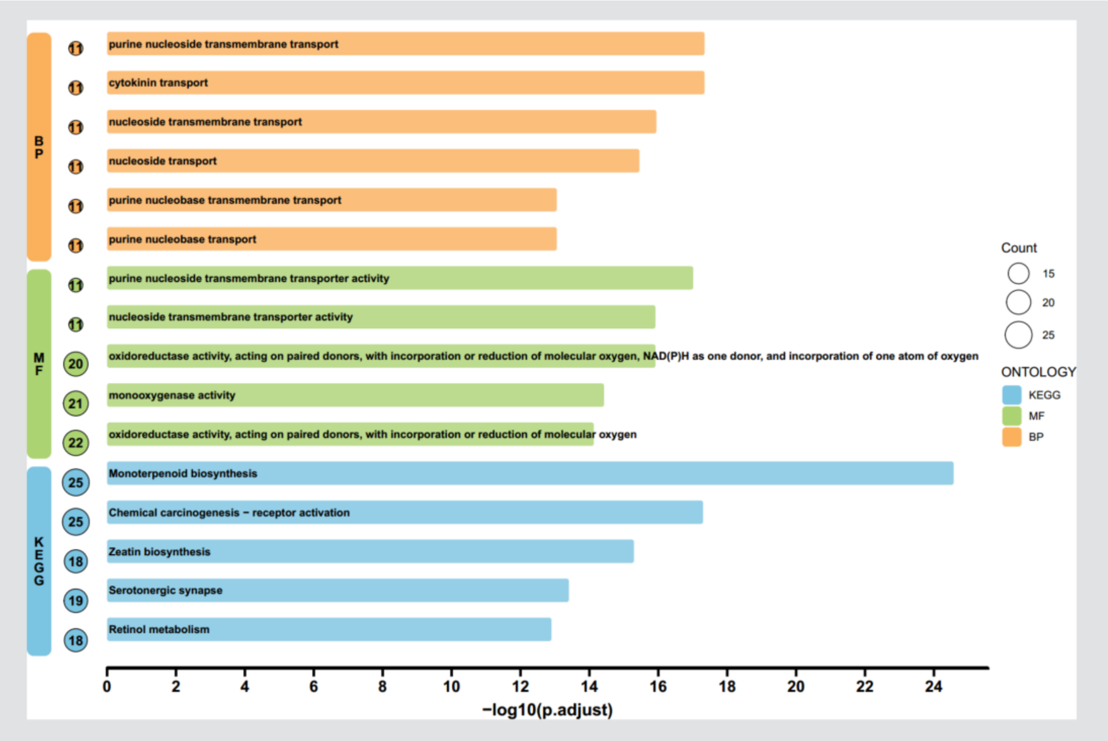


**Figure S13.** Enrichment analysis of *S. pubescens* contracted genes based on the CAFE results.

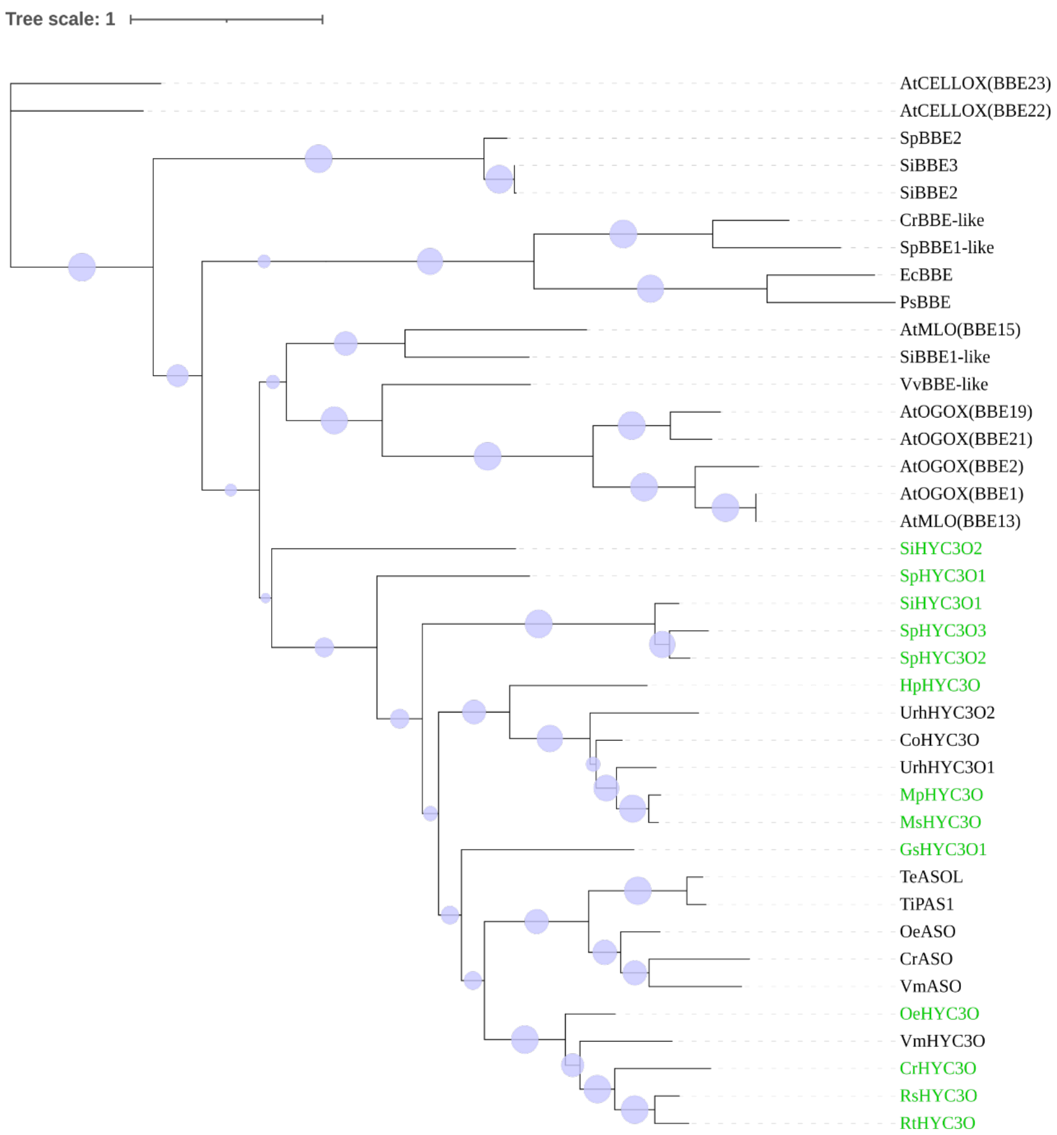


**Figure S14.** Phylogeny of MIA-oxidizing enzymes in Gentianales.

A phylogenetic tree of representative BBElike flavin-dependent oxidases was reconstructed. HYC3O (green) and O-acetylstemmadenine oxidase (ASO) sequences form a monophyletic clade within Gentianales, suggesting a single origin of these MIA-oxidizing enzymes from broader BBElike oxidase families that also include *Arabidopsis thaliana* monolignol oxidases (AtMLO) and oligosaccharide oxidases (AtOGOX and AtCELLOX). We identified HYC3O homologs in both Strychnos species (green labels), supporting the presence of an HYC3O-like lineage associated with MIA oxidation in Strychnos. All protein sequences are available from Supplementary Table14.

Abbreviations: Co, *Cephalanthus occidentalis;* Cr, *Catharanthus roseus;* Csi, *Camellia sinensis;* Gs, *Gelsemium sempervirens;* Hp, *Hamelia patens;* Hlu, *Humulus lupulus;* Ms, *Mitragyna speciosa;* Mt, *Mitragyna parvifolia;* Oe, *Ochrosia elliptica;* Rs, *Rauvolfia serpentina;* Rt, *Rauvolfia tetraphylla;* Rst, *Rhazya stricta;* Snv, *Strychnos nux-vomica;* Te, *Tabernaemontana elegans;* Ti, *Tabernanthe iboga;* Ur, *Uncaria rhynchophylla;* Vm, *Vinca minor;* At, *Arabidopsis thaliana;* Ec, *Eschscholzia californica;* Os, *Oryza sativa;* Ps, *Papaver somniferum;* Pt, *Populus tremuloides;* Ptr, *Populus trichocarpa;* Vv, *Vitis vinifera;* Si, *Strychnos ignatii;* Sp, *Strychnos pubescens.*

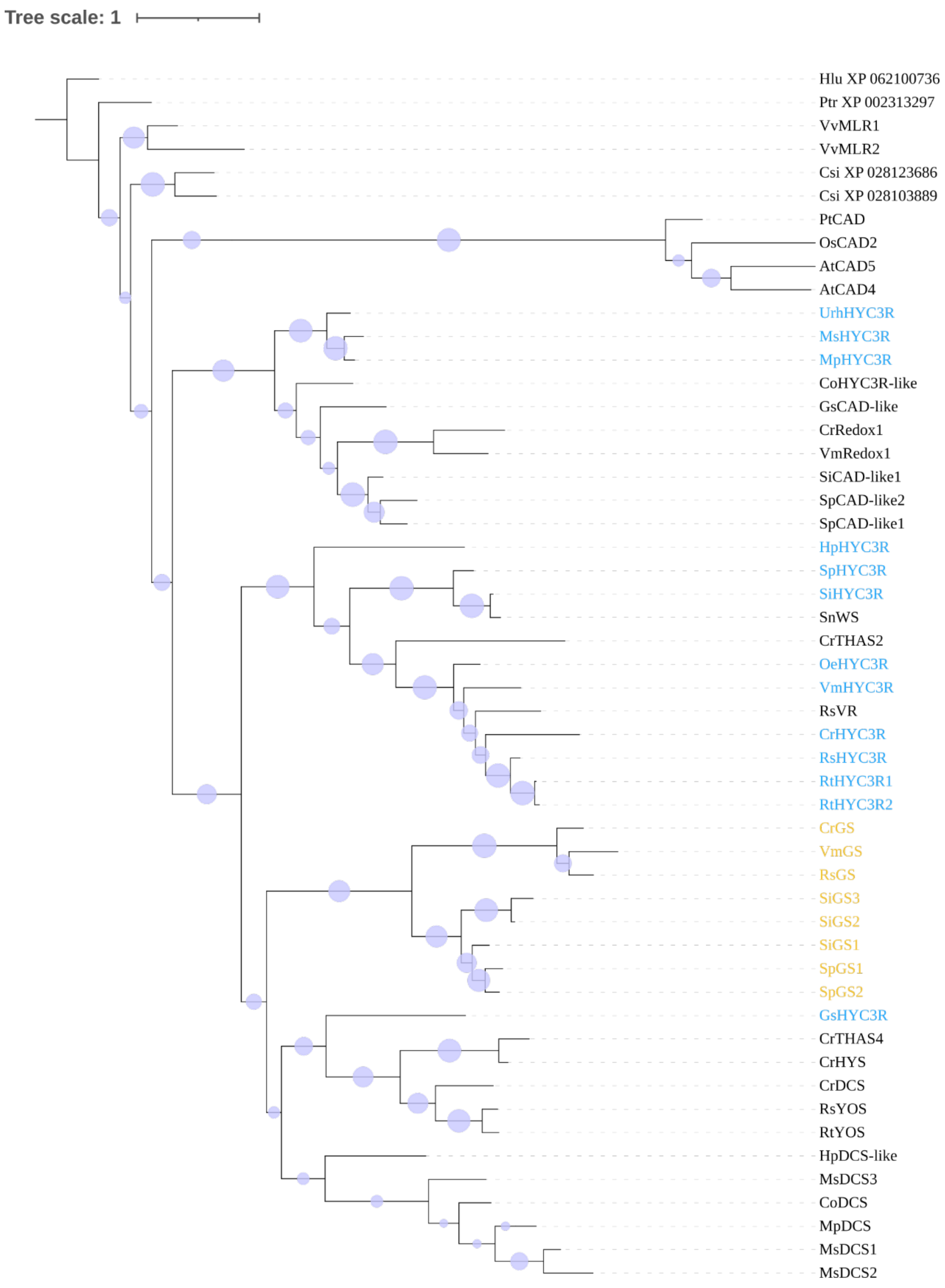


**Figure S15.** Phylogeny of HYC3R and GS enzymes in Gentianales.

Phylogenetic analysis of plant CAD-like reductases shows that MIA-reducing HYC3Rs (blue) derive from the broader family of bona fide monolignol dehydrogenases/CAD enzymes. Characterized Apocynaceae HYC3Rs together with HpHYC3R (Hamelia patens, Rubiaceae; an early-diverging Gentianales lineage) form a well-supported monophyletic clade, indicating a shared evolutionary origin. Notably, the SiHYC3R (*Strychnos ignatii*) and SpHYC3R (*Strychnos pubescens*) also fall within this HYC3R clade, supporting the presence of HYC3R homologs in Strychnos. Geissoschizine synthases (GS) are highlighted in orange, including newly identified *Strychnos* GS homologs (SiGS1–SiGS3, SpGS1–SpGS2). Node support is indicated by circle size; the scale bar represents substitutions per site. All protein sequences are available from Supplementary Table15.

Abbreviations: Co, *Cephalanthus occidentalis;* Cr, *Catharanthus roseus;* Csi, *Camellia sinensis;* Gs, *Gelsemium sempervirens;* Hp, *Hamelia patens;* Hlu, *Humulus lupulus;* Ms, *Mitragyna speciosa;* Mt, *Mitragyna parvifolia;* Oe, *Ochrosia elliptica;* Rs, *Rauvolfia serpentina;* Rt, *Rauvolfia tetraphylla;* Rst, *Rhazya stricta;* Snv, *Strychnos nux-vomica;* Te, *Tabernaemontana elegans;* Ti, *Tabernanthe iboga;* Ur, *Uncaria rhynchophylla;* Vm, *Vinca minor;* At, *Arabidopsis thaliana;* Ec, *Eschscholzia californica;* Os, *Oryza sativa;* Ps, *Papaver somniferum;* Pt, *Populus tremuloides;* Ptr, *Populus trichocarpa;* Vv, *Vitis vinifera;* Si, *Strychnos ignatii;* Sp, *Strychnos pubescens.*

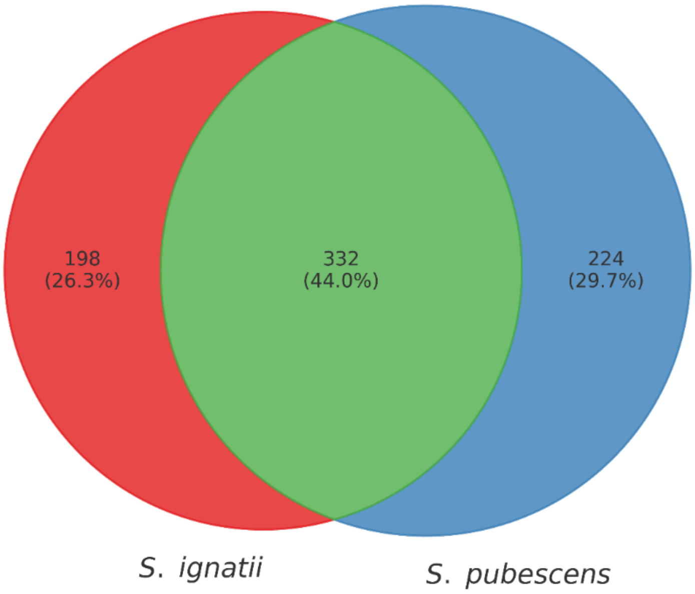


**Figure S16.** Venn plot of Terpenoid Metabolites between *S. ignatii* and *S. pubescens*

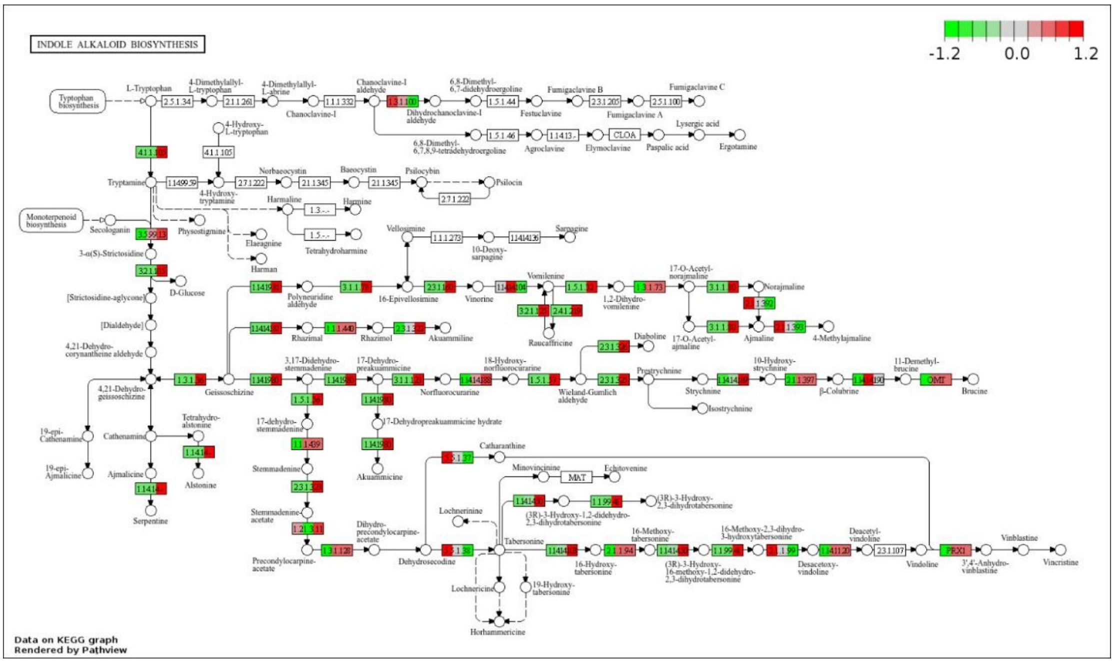


**Figure S17.** Expression profiles of homologous MIA biosynthetic pathway genes in *S. ignatii* based on transcriptome data.

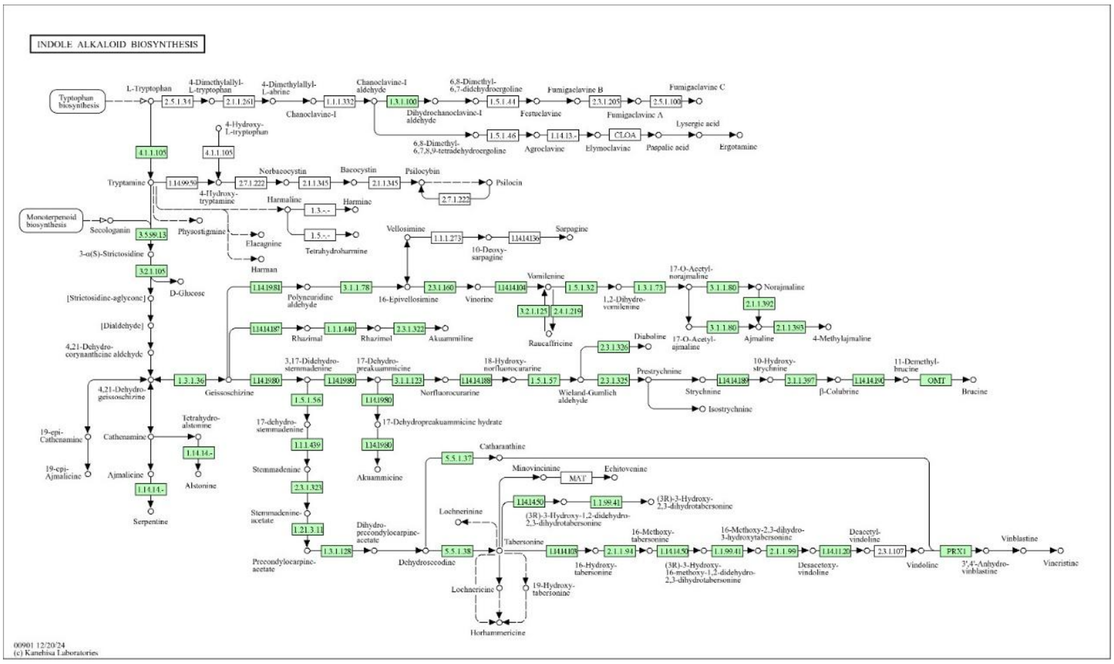


**Figure S18.** Expression profiles of homologous MIA biosynthetic pathway genes in *S. pubescens* based on transcriptome data.

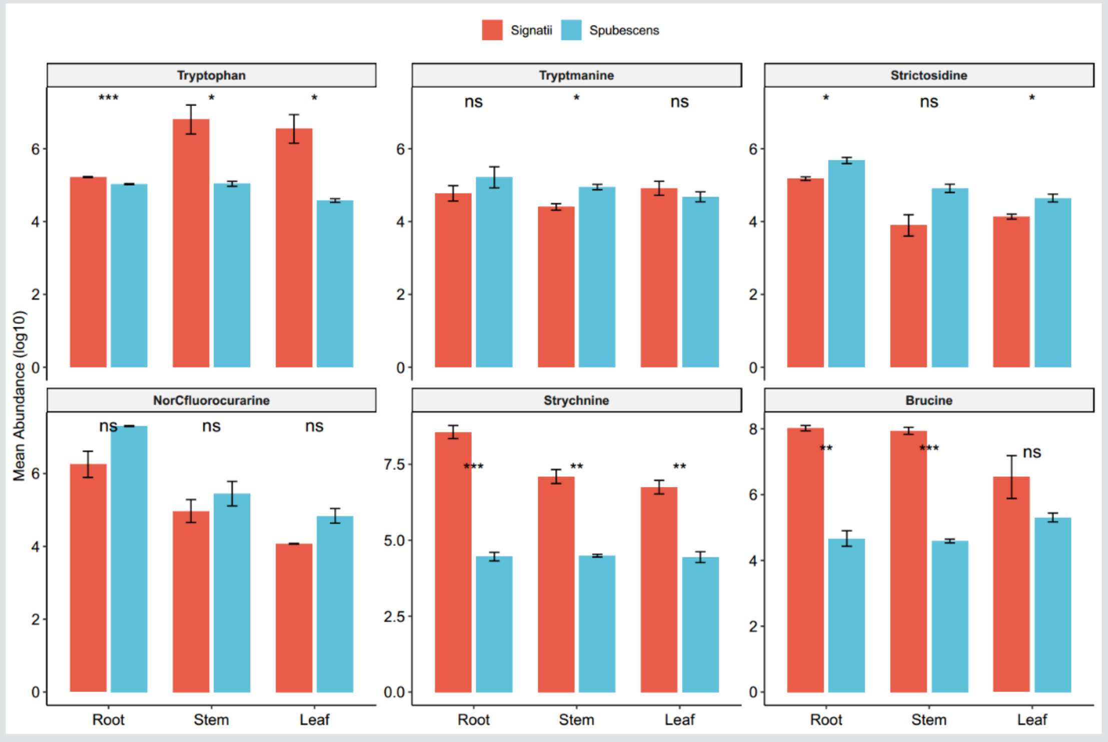


**Figure S19.** Differences in the abundances of metabolites in the strychnine biosynthetic pathway across the root, stem, and leaf tissues of *S. ignatii* and *S. pubescens*.

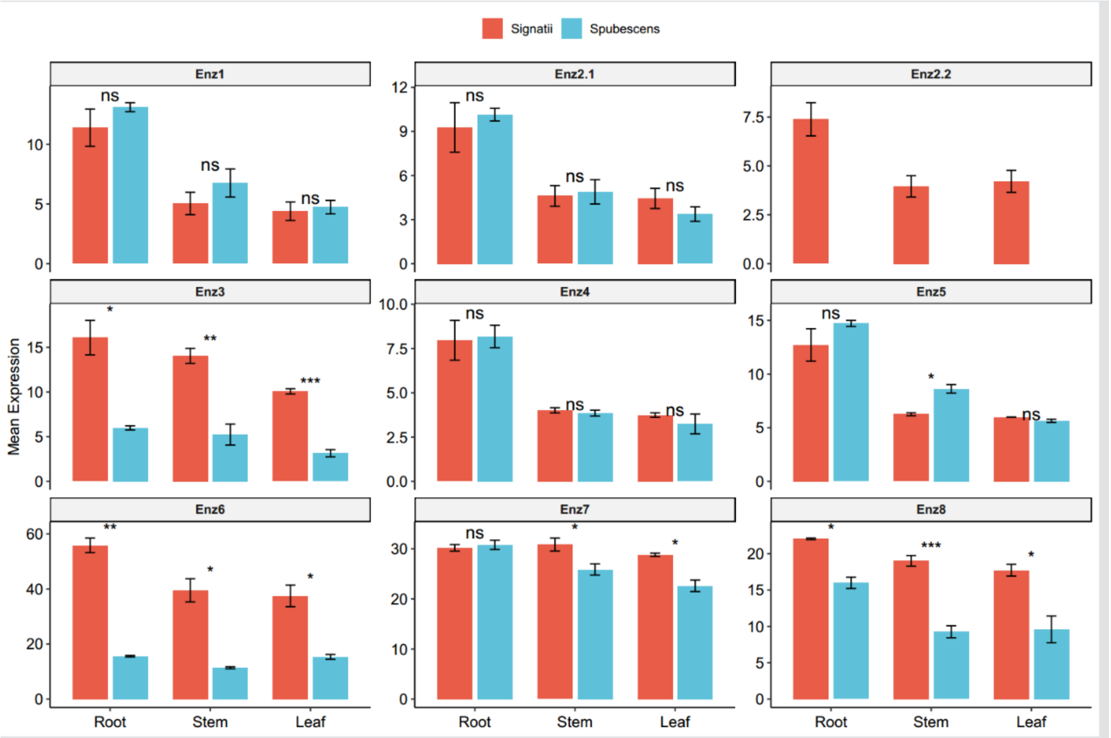


**Figure S20.** Differences in the expression levels of homologous strychnine biosynthetic pathway genes across the root, stem, and leaf tissues of *S. ignatii* and *S. pubescens*.

**Note:** Enz1 indicates geissoschizine oxidase (GO); Enz2 indicates norfluorocurarine synthase (NS); Enz3 indicates norfluorocurarine oxidase (NO); Enz4 indicates Wieland-Gumlich aldehyde synthase (WS); Enz5 indicates acetyltransferase (AT); Enz6 indicates strychnine 10-hydroxylase (10H); Enz7 indicates hydroxystrychnine O-methyltransferase (OMT); Enz8 indicates beta-colubrine 11-hydroxylase (11H).

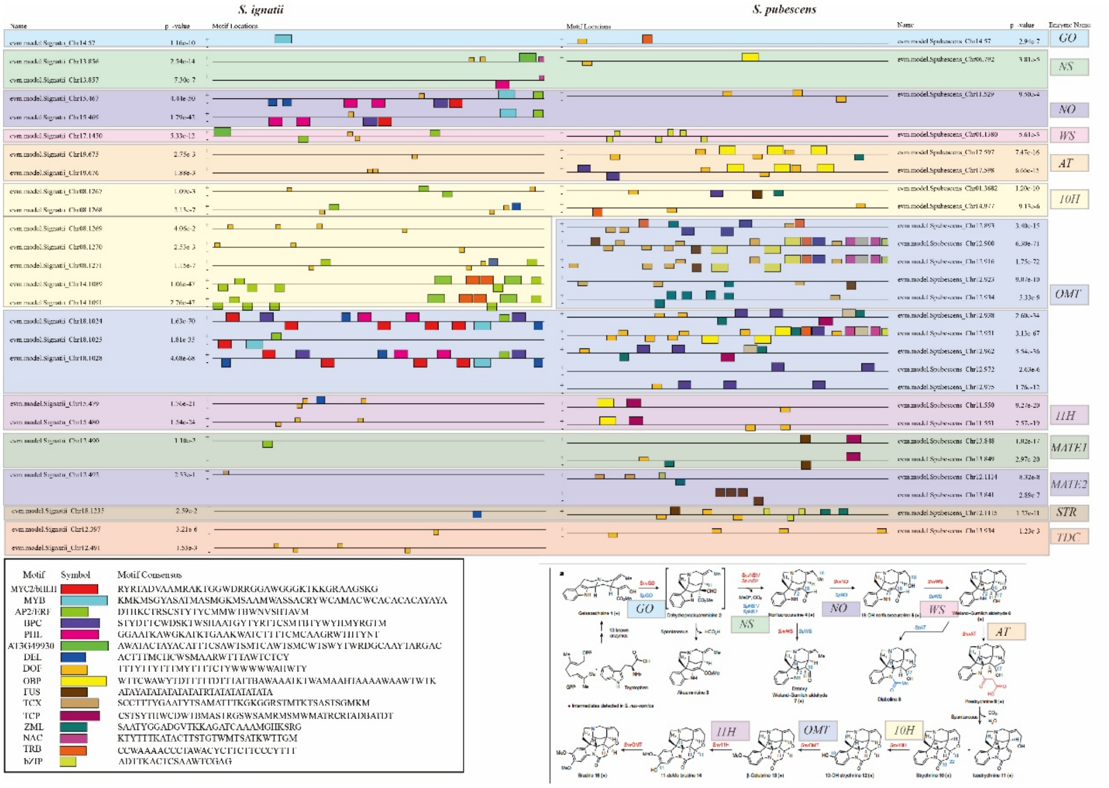

**Figure S21.** MEME-based identification of enriched motifs within the 1-kb upstream regions of MIA biosynthetic genes in *S. ignatii* and *S. pubescens* using the MEME Suite platform.

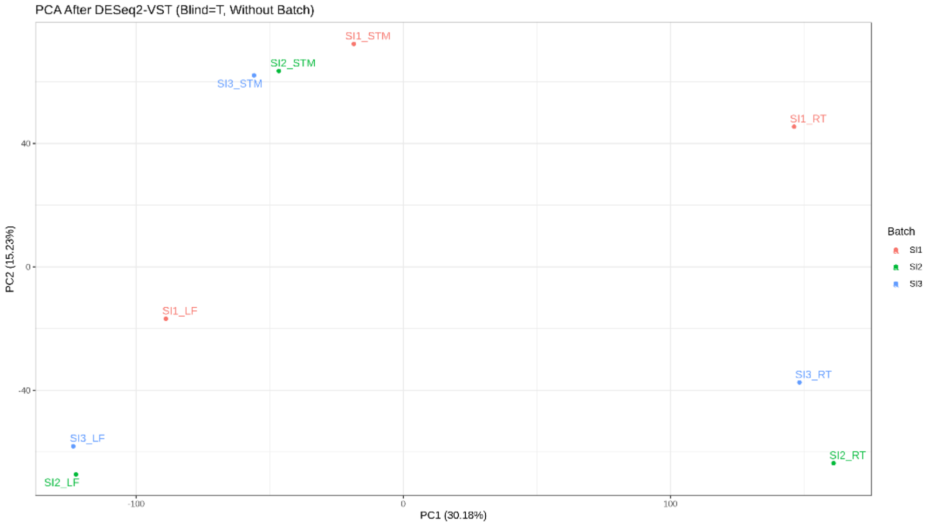


**Figure S22.** Principal component analysis (PCA) of the *S. ignatii* transcriptome.
Note: LF indicates Leaf sample; STM indicates stem sample; RT indicates root sample.

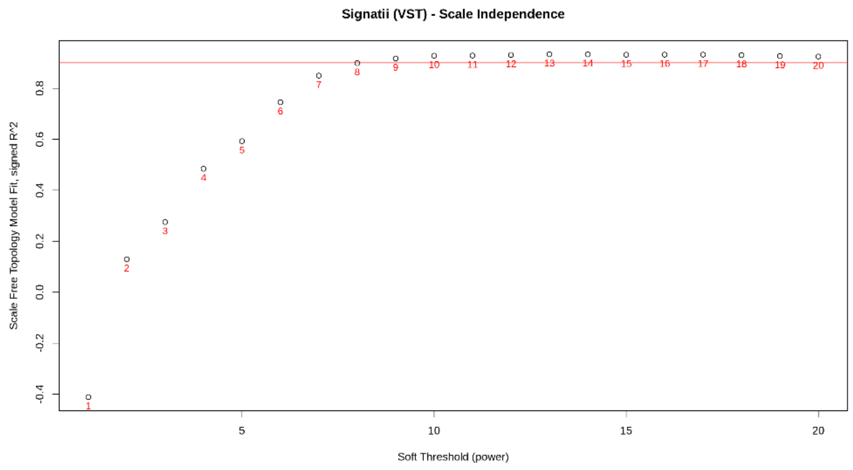


**Figure S23.** Topology model fit results of the WGCNA analysis in *S. ignatii* transcriptome.

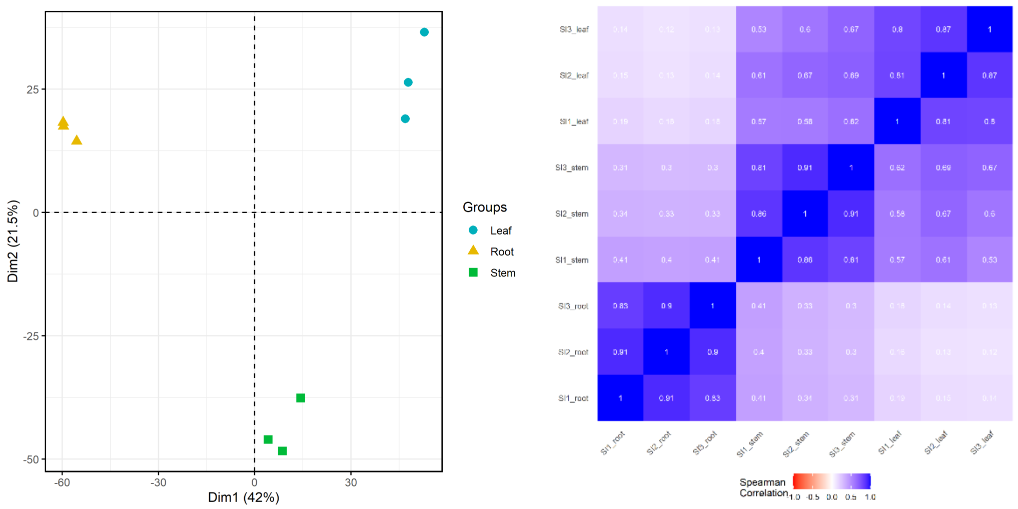


**Figure S24.** Principal component analysis (PCA) and Spearman correlation analyses of the non-targeted metabolomic profiles of *S. ignatii*.

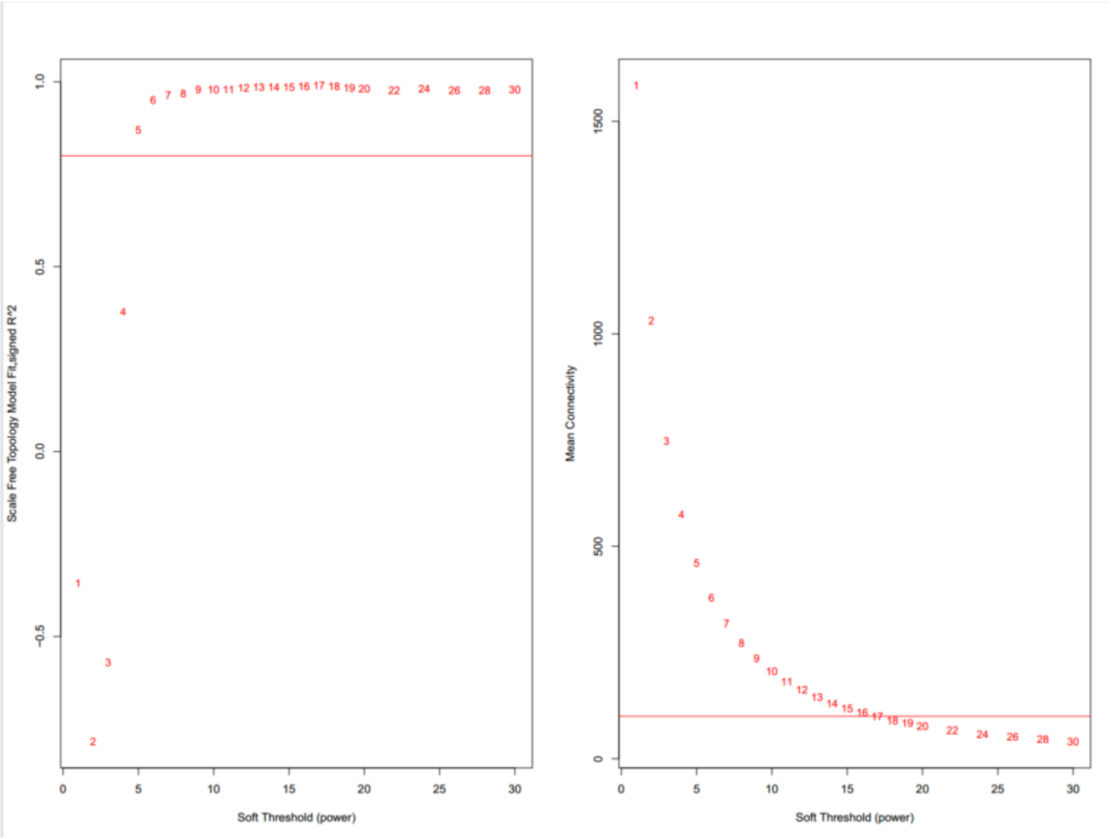


**Figure S25.** Topology model fit results of the WGCNA analysis in *S. ignatii* untargeted metabolome.

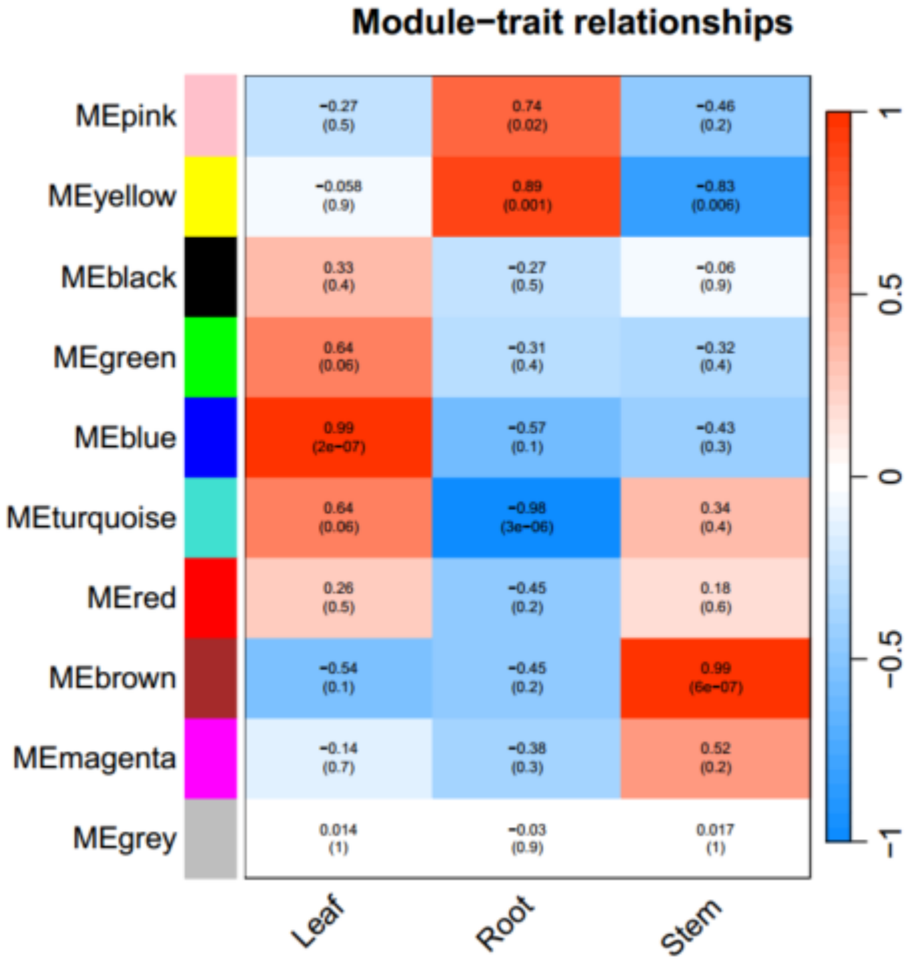


**Figure S26.** Correlation analysis of non-targeted metabolomic WGCNA modules across different organs of *S. ignatii*.

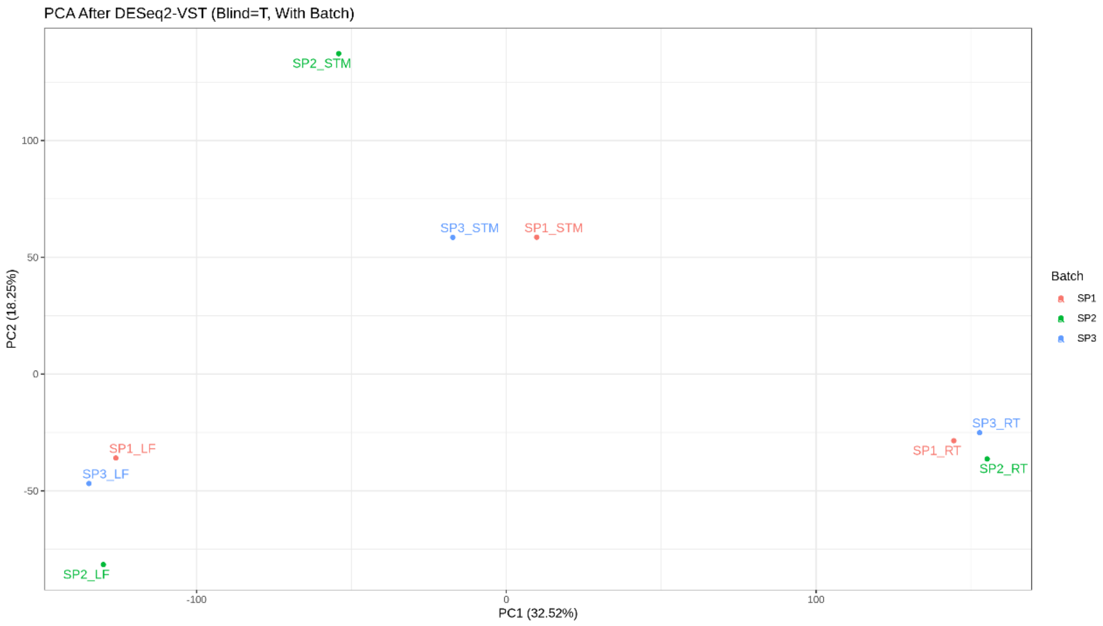


**Figure S27.** Principal component analysis (PCA) of the *S. pubescens* transcriptome. Note: LF indicates Leaf sample; STM indicates stem sample; RT indicates root sample.

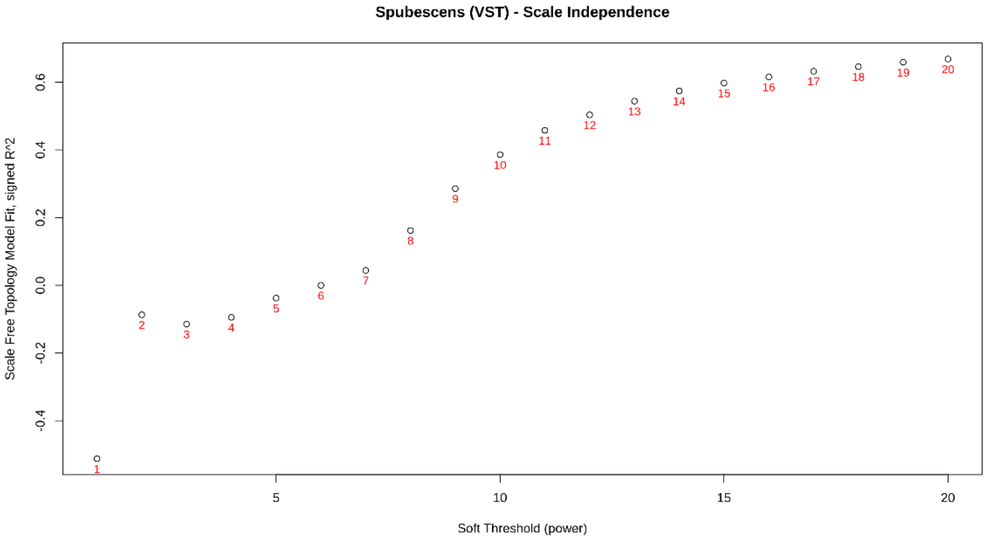


**Figure S28.** Topology model fit results of the WGCNA analysis in *S. pubescens* transcriptome.

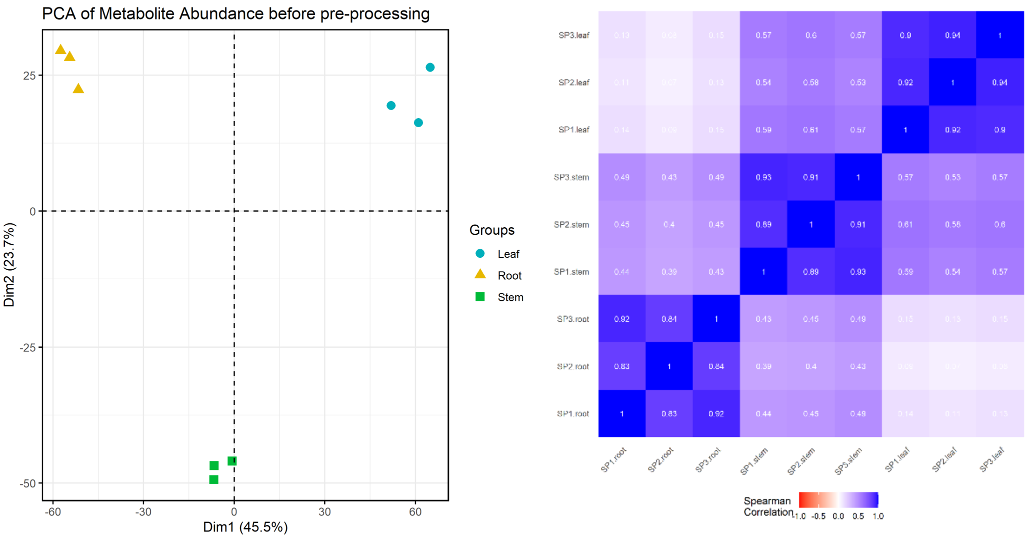


**Figure S29.** Principal component analysis (PCA) and Spearman correlation analyses of the non-targeted metabolomic profiles of *S.pubescens*.

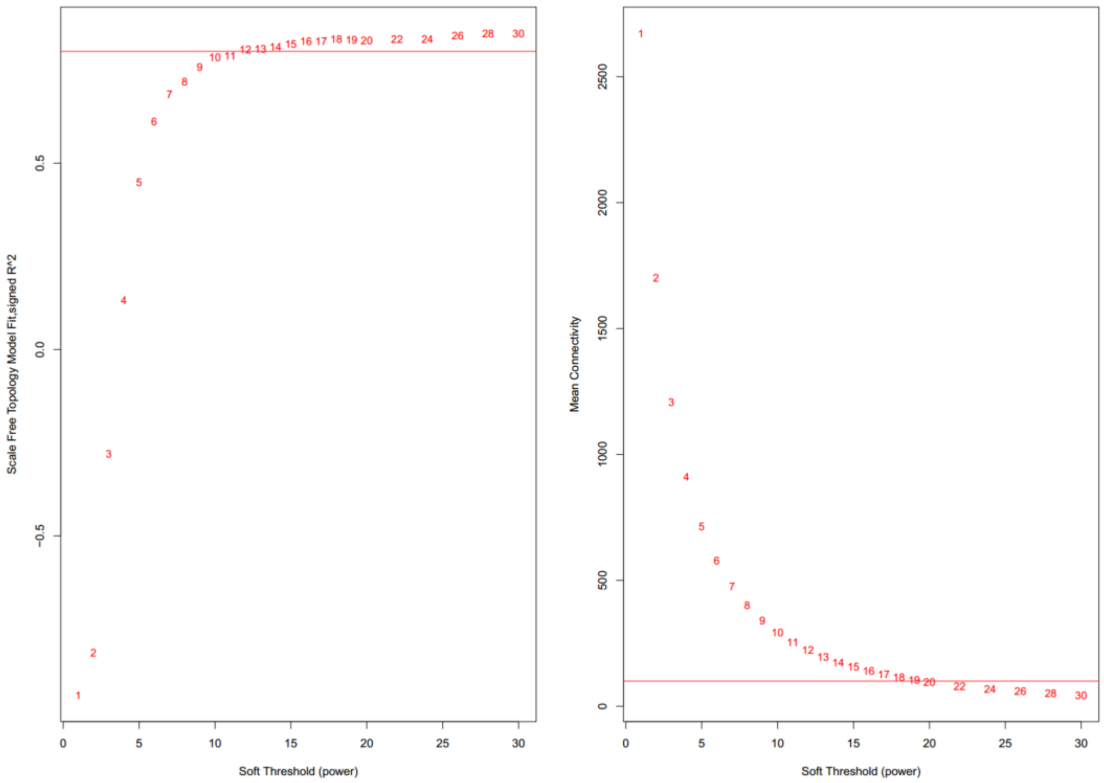


**Figure S30.** Topology model fit results of the WGCNA analysis in *S. pubescens* untargeted metabolome.

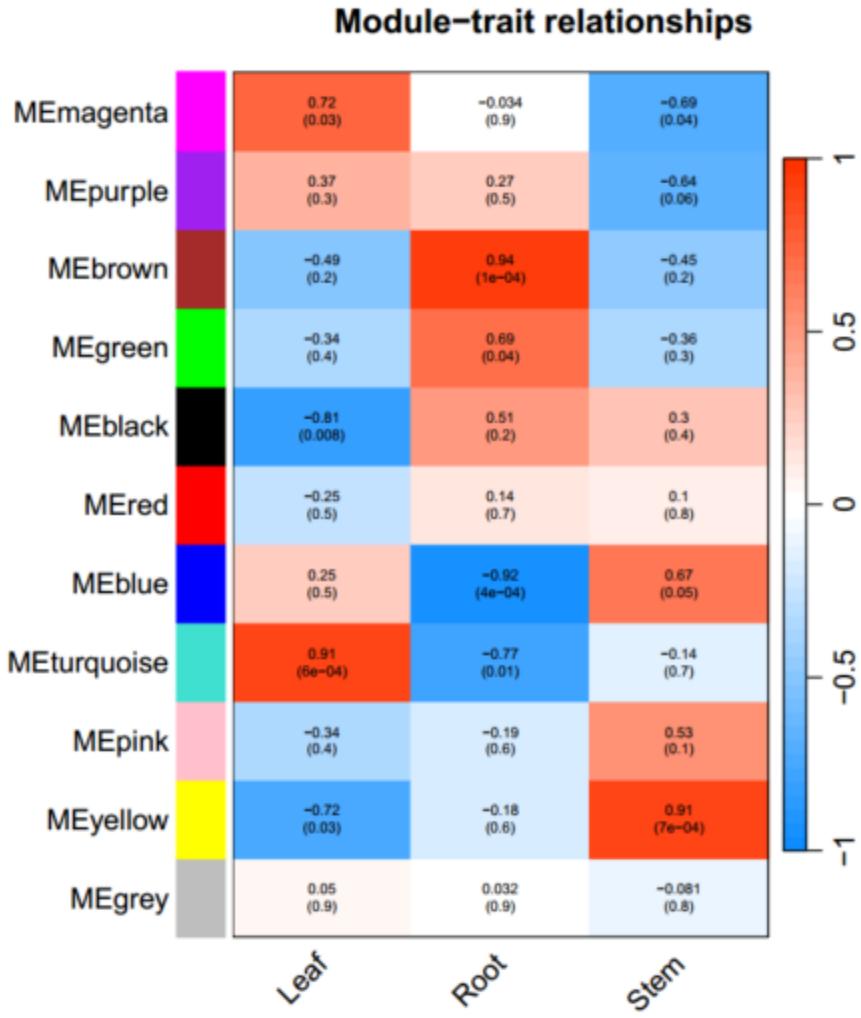


**Figure S31.** Correlation analysis of non-targeted metabolomic WGCNA modules across different organs of *S. pubescens*.

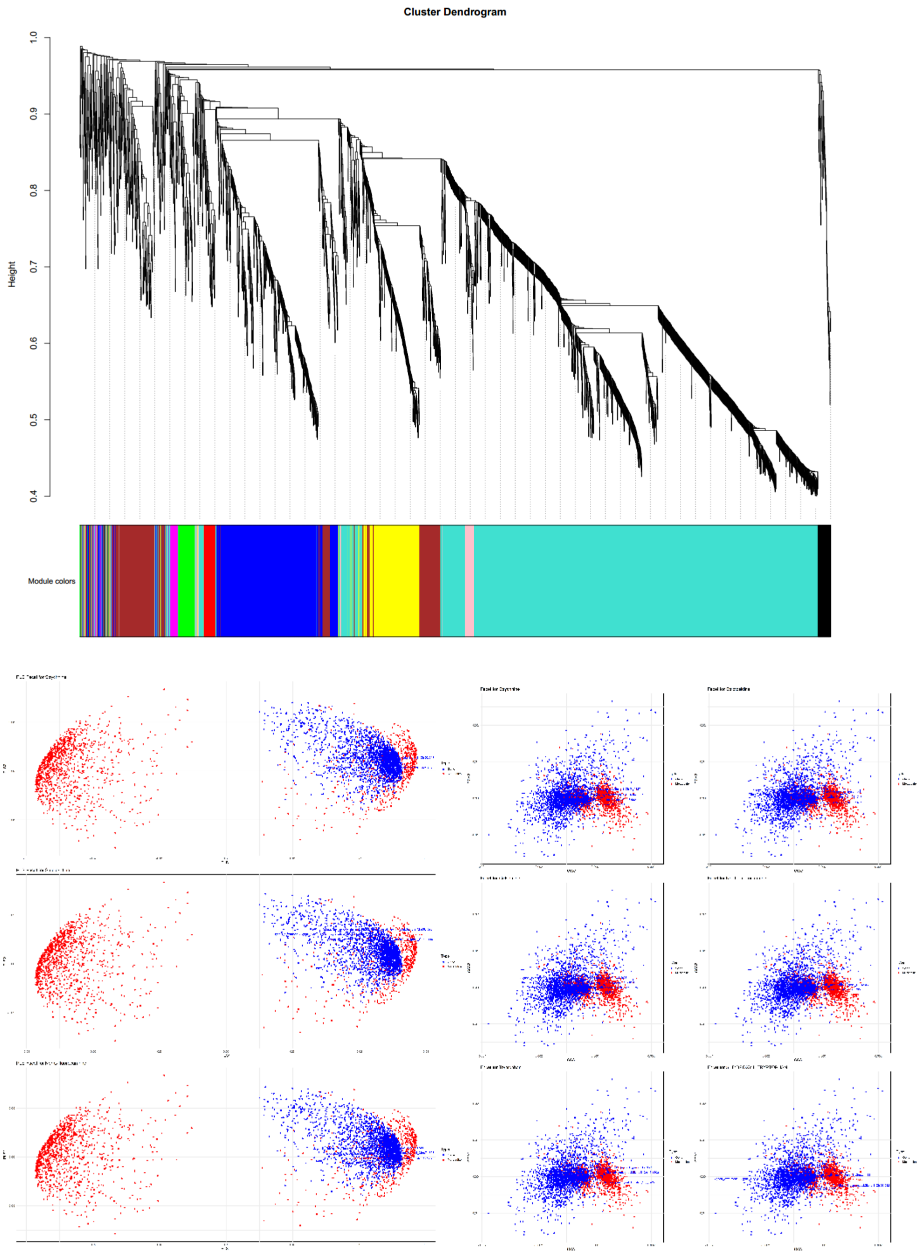


**Figure S32**. Integrative transcriptome–metabolome analyses in *S. ignatii* to identify genes associated with MIA biosynthesis.

A: WGCNA cluster dendrogram of non-targeted metabolomic profiles in *S. ignatii*. B: rCCA and PLS association scatter plot between the metabolite Turquoise module and the transcript Brown module.

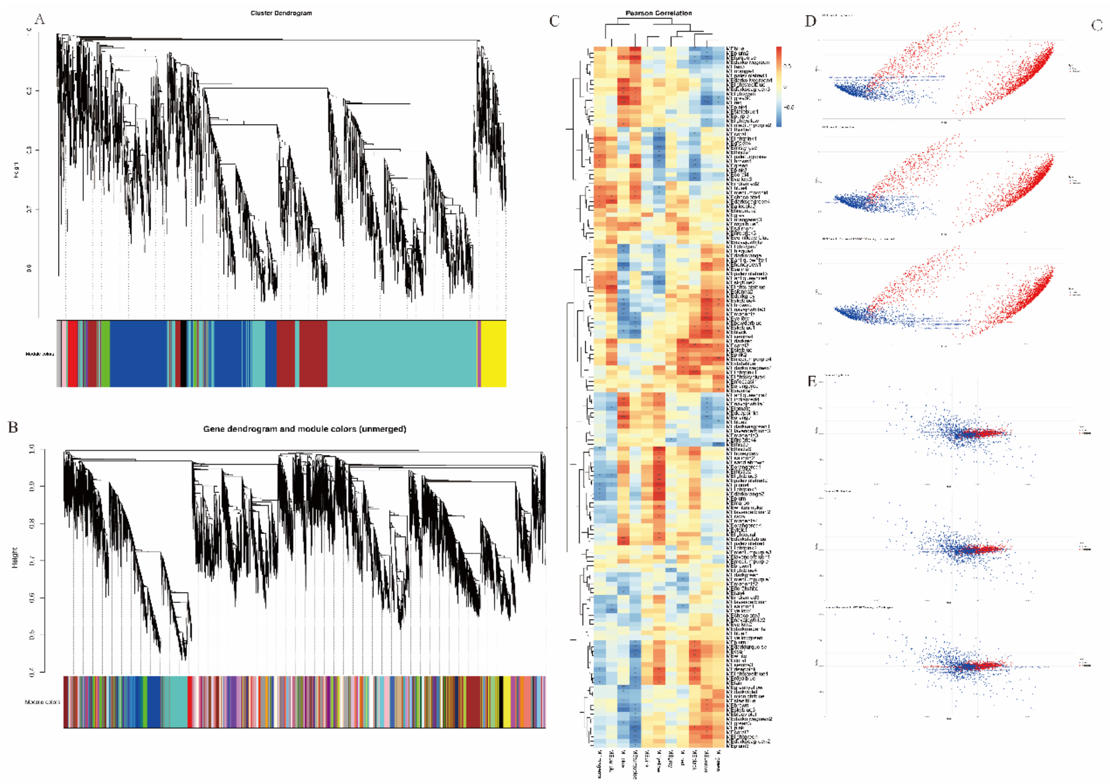


**Figure S33**. Integrative transcriptome–metabolome analyses in *S. pubescens* to identify genes associated with MIA biosynthesis.

(A) WGCNA cluster dendrogram of non-targeted metabolomic profiles in *S.pubescens*. (B) WGCNA cluster dendrogram of transcriptomic profiles in *S.pubescens*. (C) Correlation heatmap between metabolite and transcript WGCNA modules, the horizontal axis represents metabolite modules and the vertical axis represents transcript modules. (D) PLS-based association scatter plot between the metabolite Turquoise module and the transcript Yellow module. (E) rCCA-based association scatter plot between the metabolite Turquoise module and the transcript Yellow module.
